# DTFLOW: Inference and Visualization of Single-cell Pseudo-temporal Trajectories Using Diffusion Propagation

**DOI:** 10.1101/2020.09.10.290973

**Authors:** Jiangyong Wei, Tianshou Zhou, Xinan Zhang, Tianhai Tian

**Author notes:** Previous address: School of Statistics and Mathematics, Zhongnan University of Economics and Law, Wuhan, China (Jiangyong Wei). (Wei J), (Zhou T), (Zhang X), (Tian T).

## Abstract

One of the major challenges in single-cell data analysis is the determination of cellular developmental trajectories using single-cell data. Although substantial studies have been conducted in recent years, more effective methods are still strongly needed to infer the developmental processes accurately. In this work we devise a new method, named DTFLOW, for determining the pseudo-temporal trajectories with multiple branches. This method consists of two major steps: namely a new dimension reduction method (i.e. Bhattacharyya kernel feature decomposition (BKFD)) and a novel approach, named Reverse Searching on kNN Graph (RSKG), to identify the underlying multi-branching processes of cellular differentiations. In BKFD we first establish a stationary distribution for each cell to represent the transition of cellular developmental states based on the random walk with restart algorithm and then propose a new distance metric for calculating pseudo-times of single-cells by introducing the Bhattacharyya kernel matrix. The effectiveness of DTFLOW is rigorously examined by using four single-cell datasets. We compare the efficiency of the new method with two state-of-the-art methods. Simulation results suggest that our proposed method has superior accuracy and strong robustness properties for constructing pseudo-time trajectories. Availability: DTFLOW is implemented in Python and available at https://github.com/statway/DTFLOW.

## Introduction

Recent advances in single-cell technology have provided powerful tools to measure gene expression levels or protein activities of thousands single cells in one experiment. Compared with the traditional experimental studies using bulk samples that average out the responses from a large number of cells in a population, the analysis of cellular aspects at the single-cell level offers promising advantages to investigate the heterogeneity in cellular processes [1]. Since temporal data cannot be collected straightforward, a major step in single-cell studies is to order individual cells according to their progress along the differentiation pathways. The pseudo-temporal data based on the ordered individual cells will ultimately lead to the reconstruction of regulatory networks and cellular differentiation pathways [2].

Since the first algorithm Monocle for the pseudo-temporal ordering [3], a number of data-driven computational methods have been developed to define the relative position of each cell during the differentiation process. The methods for inferring pseudo-time trajectories typically consist of two major steps: namely a dimensionality reduction step and a trajectory modelling step. A class of methods based on the graph theory uses the minimum-spanning tree (MST) or shortest path to construct the major structure of trajectories, and then project all single cells onto the major structure to obtain the pseudo-time trajectory. These methods include Wanderlust [4], Wishbone [5], TSCAN [6], Monocle [3], Monocle2 [7], Waterfall [8], SCOUT [9], DensityPath [10] and SoptSC [11]. Another type of algorithms employs the probabilistic model to obtain the major structure of trajectories, such as Gpfates [12], DeLorean [13] and PhenoPath [14]. In addition, other techniques have been used to develop effective methods, include the methods based on differential equations (e.g., SCOUP [15], Pseudodynamics [16], and PBA [17]), and methods based on the principal curves such as Embeddr [18] and Slingshot [19]. Usually the algorithms based on the graph theory are more efficient, but the accuracy of the inference results is susceptible to the noise in datasets. However, methods using the probabilistic model or differential equations normally need high computational cost. Recently, a number of comparison studies have been conducted to examine the performance of these algorithms [20], and more effective methods can be found in the comprehensive review papers [21-24].

Network diffusion, also known as network propagation, has attracted much attention in recent years for identifying disease genes, genetic modules and drug targets [25]. It has also been used for manifold learning and pseudo-time calculation for single-cell data. The nonlinear dimensionality reduction algorithms based on network propagation include DCA, PHATE, etc. Among them, DCA obtains the low dimensional representation of the high-dimensional dataset by minimizing the Kullback-Leibler divergence between the observed diffusion states and parameterized-multinomial logistic distributions [26]; and PHATE generates a Markov transition matrix as the diffusion operator and then embeds the operator with the non-metric multi-dimensional scaling (MDS) approach for the visualization of single-cell datasets [27]. In addition, MAGIC alleviates the noises in single-cell datasets and learns the intrinsic biological structure and gene interaction via data diffusion [28]. Diffusion map, as a random walk approach, has also been used to explore the developmental continuum of cell-fate transitions [29, 30]. The pseudo-temporal trajectory algorithm DPT defines the diffusion pseudo-time distance between two cells using the accumulated Markov transition matrix and determines the ordering of cells based on the distances between a root cell and all other cells [31]. In fact DPT can obtain the pseudo-temporal ordering results before the dimension reduction step, and thus can detect subtle changes of gene expression.

Another important issue in single-cell studies is to identify branches in the pseudo-time trajectories in order to explore the different developmental pathways. A number of algorithms have been designed to determine the branches and optimal bifurcation points. Among them, DPT determines the branching trajectories by the correlation versus anti-correlation relationship of the dpt distances between cells [31], and Wishbone identifies two post-bifurcation cell fates using the second eigenvector of a mutual disagreement matrix [5]. In addition, SLICER uses the geodesic entropy metric for branches assignment [32]; TSCAN finds the differentiation structure based on the MST algorithm applied to the cluster centres [6]; and Monocle2 conducts the branching assignment according to the branches of the DDRtree [7]. However, the majority of these branching detection approaches can identify only one bifurcation point. More sophisticated algorithms are strongly needed to determine the branching processes with multiple bifurcation events.

In this paper, we propose a new method, named DTFLOW, for inferring the pseudo-time trajectories using single-cell data. This method uses a new manifold learning method, named Bhattacharyya kernel feature decomposition (BKFD), for the visualization of underlying dataset structure. The innovation of this algorithm includes the usage of the random walk with restart (RWR) method to transform each data point into a discrete distribution and the Bhattacharyya kernel to calculate the similarities between cells. Compared with DPT, RWR includes a free parameter that can be used to tune for better inference results. More importantly, we propose a novel distance metric based on the Bhattacharyya distance to preserve the distances along the manifold. In addition, DTFLOW uses an innovative approach named Reverse Searching on kNN Graph (RSKG) to identify the underlying multi-branching processes of cellular differentiation. The effectiveness of our proposed algorithm is rigorously examined by the application to analyse four single-cell datasets.

## Methods

This section introduces the proposed DTFLOW for the inference of psuedo-time ordering using single-cell data. **Figure 1** gives the framework of this algorithm and a brief description of the major steps. The detailed steps can be found in Supplementary Algorithm 1.

**Figure 1.**
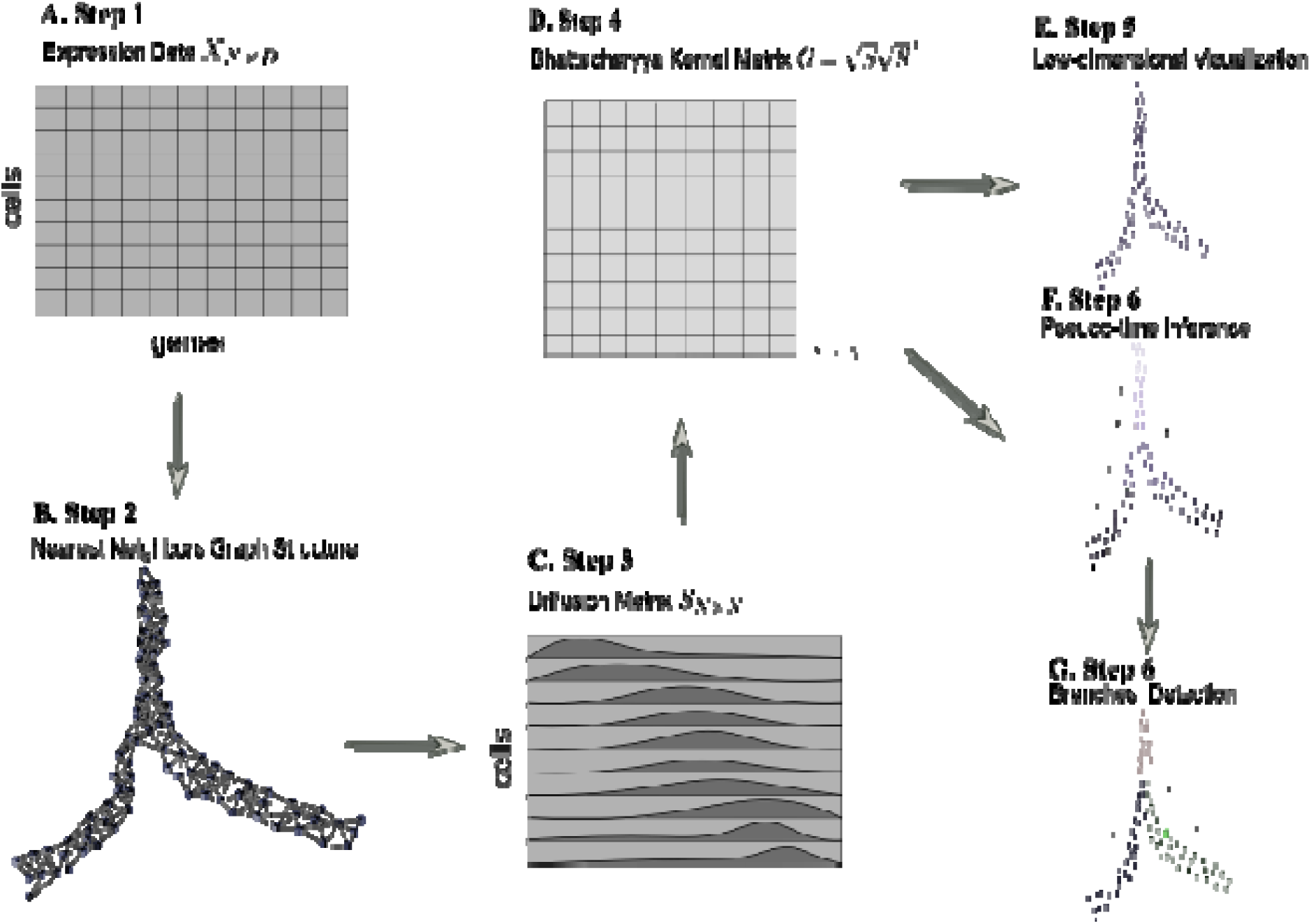
Overview of DTFLOW algorithm. A. Step 1: Pre-process of a single-cell dataset into a gene expression matrix *X*_*N*×*D*_; B. Step 2: Compute the nearest neighbours for each cell, get a nearest neighbour graph structure, and then transform the dataset into a Markov transition matrix ; C. Step 3: Using the random walk with restart method to get a diffusion matrix *S* in which each cell is represented by a discrete distribution vector;. Step 4: Construct a Bhattacharyya kernel matrix and a matrix based on the properties of kernel method; E. Step 5: Perform singular value decomposition on to get low-dimensional embedding ; F. Step 6: Calculate the new distance metric based on the row of the matrix corresponding to the root cell, and unitize it to get the pseudo-time distances ; G. Step 7: Identify the multi-branches of cellular differentiation by reverse searching based on the nearest neighbour graph structure.

### Markov Adjacency Matrix Construction

We denote *N* as the number of cells, *D* the number of genes, and *x*_*ij*_ ∈ *R*^*N*×*D*^ the gene expression data. For each cell *x*_*i*_ (i = 1, …,*N*)with expression data *{x*_*i1*_, …, *x*_*iD*_*}*, we first find its *k* most similar neighbors (include itself) through the kNN algorithm based on the pairwise cell-cell Euclidean distance. To obtain a robust diffusion operator, we construct the affinity matrix using the procedure in [33]. We first transform the cell-cell Euclidean distances into the symmetric Gaussian kernel weights to represent the affinities/similarities between cells. The transition probability between any two neighbour cells is defined by the Gaussian kernel

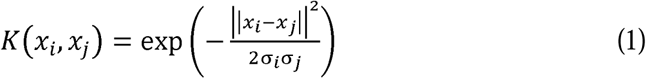

where *σ*_*i*_ and σ _*j*_ are the local kernel width of cell *x*_*i*_ and *x*_*j*_, respectively. The value of *σ*_*i*_ is based on the local density with its distance to the *k*-th nearest neighbour. If cell *x*_*i*_ is a neighbour of *x*_*j*_ but *x*_*j*_ is not a neighbour of *x*_*i*_, we let *K(x*_*j*_,*x*_*i*_*) = K(x*_*i*_,*x*_*j*_) to generate a symmetric kernel matrix. If *x*_*i*_ and *x*_*j*_ are not the neighbour of each other, *K(x*_*i*_,*x*_*j*_) = 0 Then we normalize the kernel as

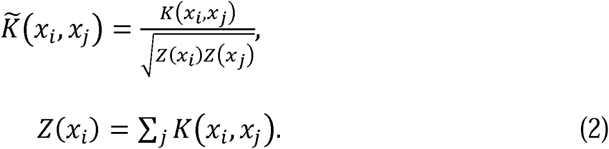

Finally, we define the Markov transition probability matrix using the normalization over rows, defined by

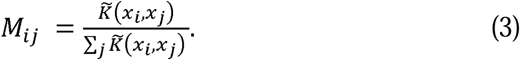

### Bhattacharyya kernel feature decomposition

Random walk with restart (RWR), also known as Personalized PageRank (PPR) [34], is a ranking algorithm that has a number of good properties such as the ability to capture the global structure of a graph and multi-facet relationship between two nodes in the graph [35]. It has a wide range of applications in link prediction, community detection, and anomaly detection. In RWR, each cell is considered as a node, and RWR iteratively calculates the relevance (proximity) score of each node with regard to a given seed node *i* in the kNN graph. At each step *t*, RWR select a move from the current node either to its neighbours with probability *p*, or return to itself with the restart probability 1 − *p*. Then the distribution vector satisfies the following equation:

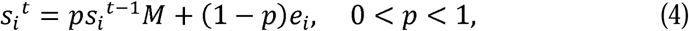

where *s*_*i*_^*t*^ is an *N*-dimensional row distribution vector for the visiting probability of each node from the seed node *i* after t steps, *M* is the transition matrix defined by (3), and *s*_*i*_^*0*^ *= e*_*i*_ is a unit direction row vector, which means that the propagation starts from node *i*. Thus RWR can be regarded as a more general approach and DPT is a special case of RWR (i.e. *p* = 1) (see Supplemental Information). We introduce the stationary distribution for the visiting probabilities by iterating the updating step (4) infinitely (i.e. *t* → ∞) until the convergence of the distribution vector. Then the final stationary distribution is

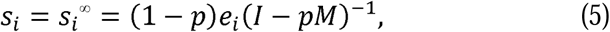

where *I* is the identity matrix. The diffusion matrix *S =* [*s*_1_, …, *S*_*N*_]^*T*^ is writen as

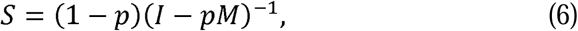

The diffusion distribution of each node is a non-vanishing distribution, i.e. *s*_*ij*_ *>* 0 and 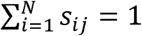, where element *s*_*ij*_ of matrix *S* is the similarity score of node *j* towards node *i*. Cells with a smaller similarity score should have ordering that is farther away than those having larger similarity scores. Based on the properties of RWR, it is assumed that the exact ordering of all nodes is determined by the diffusion process, and each node is represented by a discrete distribution vector in the matrix *S*.

Suppose that *p* and *q* are two discrete probability distributions over the same space Ω = *{x*_*1*_, …, *x* _*N*_*}*, and let *p*_*i*_ *= p(x*_*i*_) and *q*_*i*_ *= q(x*_*i*_). Then Bhattacharyya coefficient (BC) measures the similarity between *p* and *q*, given by

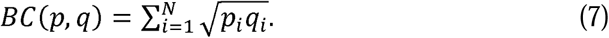

Based on the above definition, the Bhattacharyya kernel matrix [36] is defined by

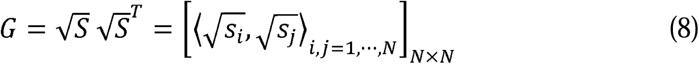

where the square root operation is conducted for every element of the matrix, and ⟨ ·,· ⟩ is the inner product of two vectors. Apparently the diagonal element of matrix *G* is the inner product of vector 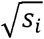 and has the value of unit one. Because *G* is a kernel matrix, its eigenvalues are greater than or equal to 0.

According to Mercer’s Theorem, there exists a kernel function *k*, satisfying that

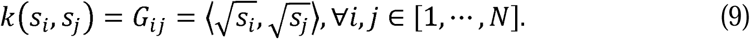

Based on the properties of kernel functions, we construct a new kernel *k*_1_ with the mapping operator *ϕ*, defined by

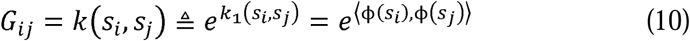

Let *y*_*i*_ *= ϕ(s*_*i*_), the above equation can be written as ⟨ *y*_*i*_,*y*_*j*_ ⟩ *=* log*G*_*ij*_,. Then we rewrite it in the matrix form

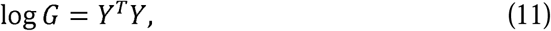

where *Y = [y*_*1*_,*···, y*_*N*_]^*T*^, and the where logarithm operation is applied to every element of *G*. Equation (11) is a linear matrix transformation, and we perform SVD to obtain decomposition

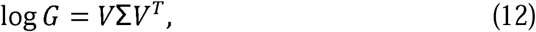

*V* ∈ *R*^*N*×*N*^ is an unitary matrix which satisfies *V*^*T*^*V = I*, and Σ is a diagonal matrix whose elements are the singular values of matrix log*G*. For the purpose of visualization, we use the largest *d* (positive) singular values to represent the major feature of matrix log*G*. The *d* low-dimensional embedding of *Y*, defined by

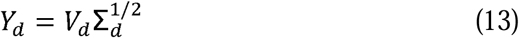

is used to represent the single-cell dataset. Here ∑_*d*_ is a matrix that includes only the largest *d* singular values and *V*_*d*_ is the corresponding vectors. Normally we use *d* = 2 or *d* = 3 for 2-dimensional or 3-dimensional visualization. Then we use the low-dimensional dataset *Y*_*d*_ to visualize the underlying structure of the original high-dimensional single-cell dataset. In this way we design a novel method for dimensionality reduction by introducing the new Bhattacharyya kernel function.

### Pseudo-time Ordering

Note that the Bhattacharyya distance [37], defined by

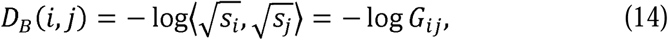

is a measure of similarity between two probability distributions. Although it has been widely used in engineering and statistical sciences, this metric is a measure of divergence, and does not satisfy the triangle inequality in the inner product space. Thus, it is not appropriate to use this metric to calculate the distances between cells. In this work, we introduce a new distance metric to measure the distance between two cells. From equation (11), we obtain the distance of two cells *i* and *j* as

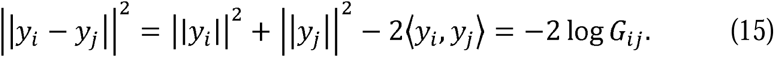

Since 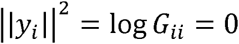, we define the new distance metric as

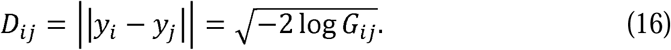

It can be shown that this new distance satisfies the triangle inequality in the inner product space, which is essentially a kernel distance [38], and this distance metric is defined on the manifold. Thus, we propose to use this metric to calculate the distance between cells.

If the root cell *x*_*r*_ is known, the distance between the root cell and the i-th cell is denoted as *D*_*ri*_, and we use the vector *T*_*r*_ *= D*_*r*,:_. to denote the pseudo-times of single cells. However, if we do not know the root cell in advance, we can select a group of cells as the root cells based on the sum of distances between a particular cell and all other cells. Suppose we select *R* cells as the group of root cells, the pseudo-times of single cells is given by 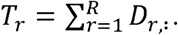. Finally we normalize the pseudo-times to values between 0 and 1, given by

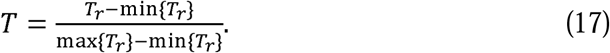

### Reverse-searching in kNN graph for branch detection

According to the Waddington’s epigenetic landscape model [39], single cells will not change their fates after differentiation. Based on the constructed kNN graph and pseudo-time of each cell, we next propose a new method for branching detection using the reverse-searching in kNN graph (RSKG). Supplementary Figure S1 shows a brief description of RSKG for identifying multi-branching processes. In this algorithm, *n* is the minimum number of cells required for forming one sub-branch, *T* the set of pseudo-times of all cells, and *A* the set of indices array of the kNN graph of all cells. For the *id*-th cell, *A* [*id*] is the set of its *k*-nearest neighbours. In addition, we use to store the reverse index ordering based on. We also use a nested list *prop-groups* to store the candidate sub-branches/groups and a nested list *sub-branches* to store the determined sub-branches. Initially these two nested lists are empty. The major steps of this algorithm are described in Supplementary Algorithm 2 together with Figure S1 (see supplemental materials).

This algorithm starts from the cell with the largest pseudo-time, whose index is the first element in. We put the indexes of its neighbour in the nested list *prop-groups* as the first candidate group. Then we consider the next element in and its neighbour. If set has intersections with the list, then extend to the list in *prop-groups*. Otherwise, append the list to *prop-groups* as a separate group. The similar procedure is applied to the following elements with index.

For the following cells, if has intersections with two or more candidate groups in *prop-groups*, and if the length of two or more intersected lists reaches, these lists will be moved from *prop-groups* to *sub-branches* and become a determined branch; otherwise, if the length of the merged list does not reach *n*, merge these lists together as one new list in *prop-groups*. If has intersections with lists in both *prop-groups* and *sub-branches*, and if the length of and the intersected list in *prop-groups* reaches, and the intersected list will be moved from *prop-groups* to *sub-branches* to become a determined branch; otherwise, the elements of and the intersected list in*prop-groups* will be assigned to the branches in *sub-branches* that are closer to them.

Our method ensures that all the end points of sub-branches can be connected within the kNN graph, and all pseudo-times of single cells in the previous sub-branch are less than those in the subsequent sub-branch(es) along the development process. Thus this new algorithm can provide more accurate inference results than the existing methods.

## Results

In this section, we evaluate the robustness and accuracy of our proposed algorithm DTFLOW for the inference of psuedo-time ordering using single-cell data. We apply DTFLOW to analyse four real scRNA-seq datasets, namely the mouse embryo dataset [40], myeloid progenitor MARS-seq dataset [41], female gonad scRNA-seq dataset [42], and microwell-seq data [43]. This work does not include any work for the pre-processing of experimental data. We use the datasets with the same input (namely the same genes and same single-cells) from the published papers directly.

### Mouse embryo single-cell dataset

The high-throughput reverse transcription polymerase chain reaction (RT-PCR) dataset [40] describes the early-stages of the developmental process for mouse embryo. This dataset includes the expression levels of 48 selected genes in 438 single cells at seven different developmental stages, namely from the 1-cell zygote stage to the 64-cell blastocyst stage. We first apply DTFLOW to project the 48-dimensional gene expression data into the two-dimensional feature space by using the BKFD algorithm. **Figure 2A** provides the visualization of single cells at different stages. It clearly reveals the seven developmental stages/labels (i.e. 1-cell stage, 2-cell stage, …, 64-cell stage), which also validates the effectiveness of our proposed dimensional reduction technique. Since not knowing the root cell in the dataset, we select a cell in the initial time stage, which has the largest sum of distances to all other cells, as the root cell. The differentiation process of single cells is characterized by the calculated pseudo-times in **Figure 2B**, whose values range from zero of the first cell to the maximal value of unit one of the last cell. These results suggest that the pseudo-times of individual cells are recovered successfully.

**Figure 2.**
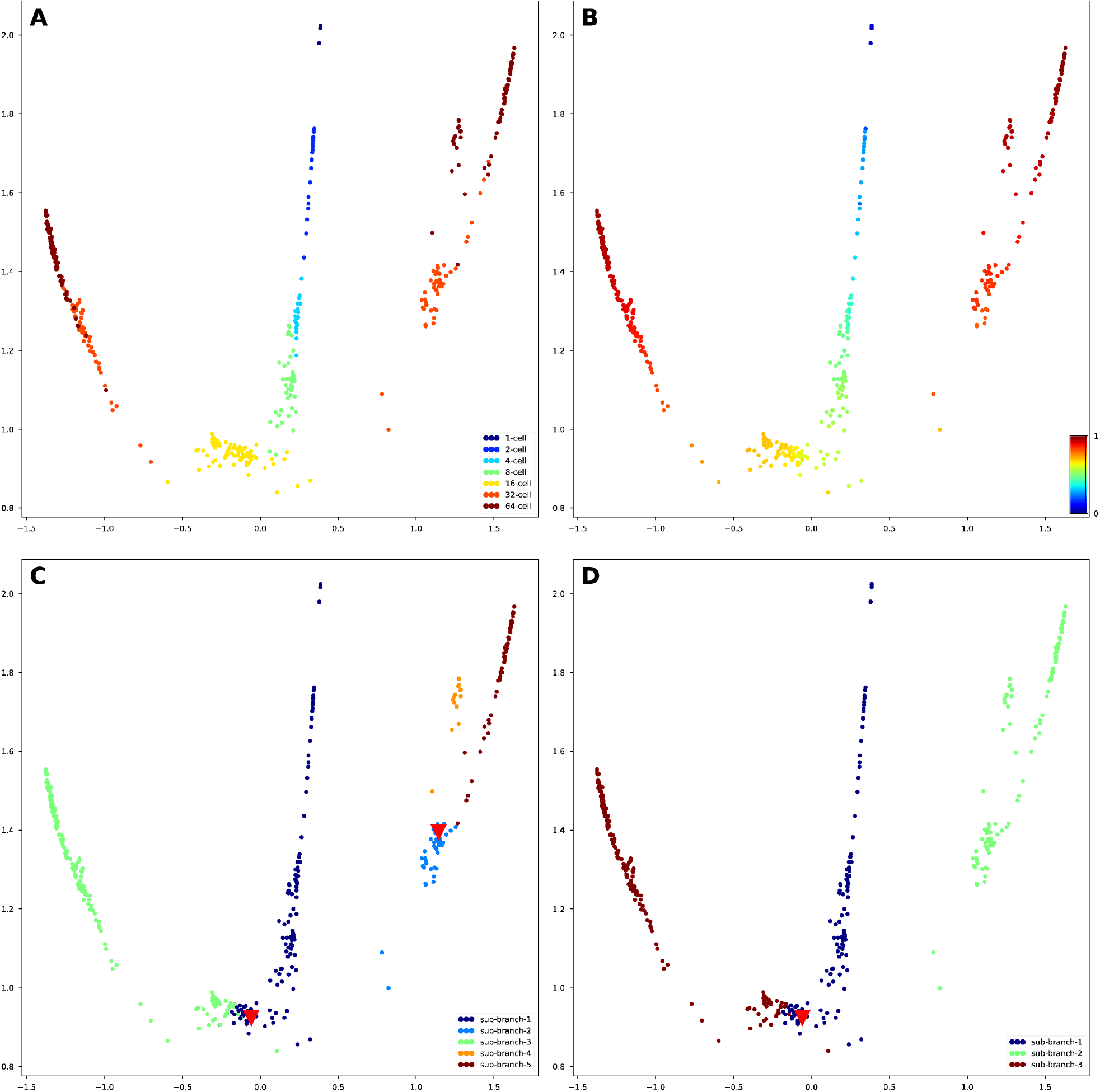
Developmental trajectories inferred by DTFLOW for the mouse embryo single-cell dataset. A. Visualization of different time stages. B. Visualization of inferred temporal trajectory. C. Visualization of inferred 5 sub-branches when the minimal cell number required for forming a sub-branch satisfies. D. Visualization of inferred 3 sub-branches when the minimal cell number required for forming a sub-branch is larger ().

We also test the influence of the minimal cell number *n* required for forming a sub-branch. When we set a small value (i.e. *n ≤* 11), the individual cells in the lineage process is divided into 5 sub-branches in **Figure 2C**. There are two bifurcation points that separate cells into two distinct sub-branches along the differentiation process. **Figure 2C** shows that the main lineage trajectory contains two major branches and one of them further differentiates into two smaller branches. It also suggests that cell differentiation does not occur in the early stages, but cells in the 32-cell stage differentiate distinctly into trophectoderm (TE) and inner cell mass (ICM). Subsequently, cells in the ICM stage further differentiate into epiblast (EPI) and primitive endoderm (PE) in the 64-cell stage. After the second bifurcating event, the embryo cells are divided into three distinct types: namely TE, PE, and EPI. However, if we use a relatively large value of *n* (= 12 ∼ 112), the single cells will form only three sub-branches with the first bifurcation event occurred in **Figure 2D**. The second bifurcation event is not identified since the lengths of sub-branches are less than the minimal cell number *n*. We use red triangles in **Figures 2C** and **2D** to indicate the bifurcation points. Note that the distances between cells in our algorithm are calculated based on the high-dimensional Bhattacharyya kernel matrix. However, the data visualized in **Figure 2** are the low-dimensional data after the application of SVD.

In our proposed algorithm, there are two free parameters that should be determined based on the datasets it applied. The first one is the number of closest neighbours *k* of each data point, which is taken into account for the determination of affinity with classes. To place greater emphasis on the local properties of the manifold structure, a smaller value of *k* is preferred. Meanwhile, the value of *k* should also be large enough for the connectivity of the kNN graph. The value of *k* in BKFD is usually smaller than that in diffusion maps for dimension reduction, which implies that BKFD can capture the local structure of manifold better than diffusion maps. We test different values of *k* and find that the results are better if *k* = 10, which will be used in this work for analyzing other datasets. The second parameter is the restart probability 1 − *p* that controls the relative influence of both local and global topological structure. To smooth the noise of data, a larger value of *p* (i.e. a smaller value of 1 − *p)* may be preferred. To test the influence of *p*, we calculate the pseudo-ordering of single cells using different values of *p*. We use the Kendall rank correlation coefficient of the inference results to compare the accuracy of the algorithms. Since knowing the stage number of each cell in the experimental data, we determine the stage number of each cell in the inferred trajectories and then calculate the Kendall rank correlation coefficient of these two types of stage numbers. An algorithm has better accuracy if the value of this correlation coefficient is larger. Numerical results in **Figure 3A** suggest that the ordering accuracy is better when the value of *p* is around 0.9. Thus, we use *p =* 0.9 in this work, including the results shown in **Figure 2**.

**Figure 3.**
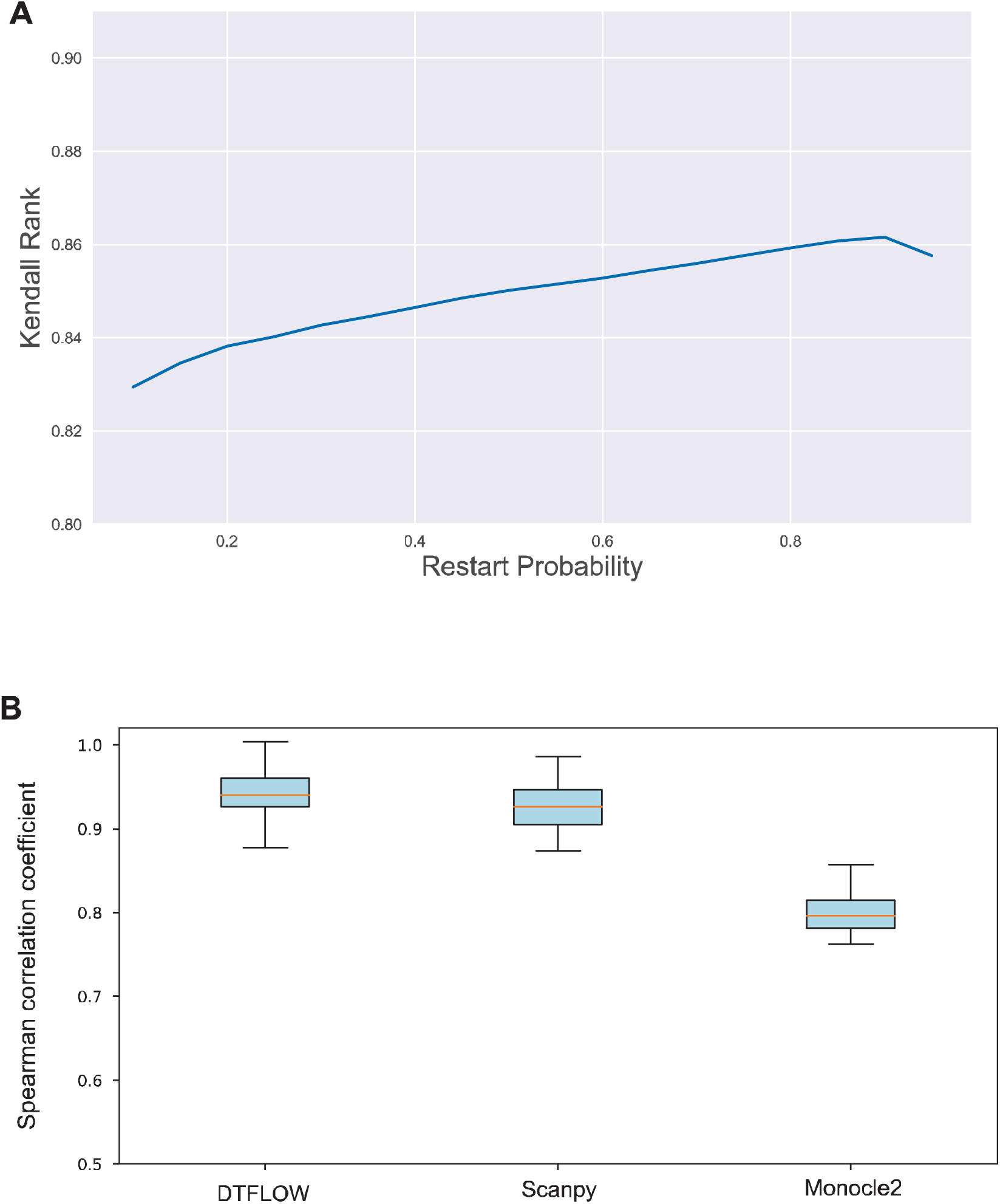
Performance of three inference methods for the mouse embryo single-cell dataset. A. The accuracy of DTFLOW that is measured by Kendall rank correlation coefficient for the networks determined by different values of restart probability. B. Robustness properties of three inference methods. The properties are obtained by randomly sampling 90% of single cells from the whole dataset and comparing the ordering results of the subset with those of the whole dataset.

To demonstrate the effectiveness of our proposed algorithm, we compare the performance of DTFLOW with two published state-of-the-art methods, namely DPT and Monocle2 [7]. We use the Python toolkit Scanpy [44] for the implementation of DPT. Supplementary Figures S2C and S3C show the branching detection results of Scanpy and Monocle2, respectively. Figure S2C suggests that Scanpy detects only three groups/sub-branches. It fails to identify the number of terminal states correctly, and also obtains the wrong location of bifurcation point. Although Monocle2 identifies three types of the terminal cells correctly in Figure S3C, it does not reveal the intermediate state between state 1 and states 3/4 (i.e. the ICM stage) using the dimensional reduction method DDRTree. For this dataset, we use Kendall rank correlation coefficient to compare the accuracy of these three algorithms. The calculated Kendall rank correlations are 0.862, 0.796, 0.761 for DTFLOW, Scanpy and Monocle2, respectively, which suggests that our proposed method has better accuracy than the two published methods.

Supplementary Figure S4 shows the expression levels of two genes, namely *Gata3* and *Sox2*, based on the inferred pseudo-times using the three methods, which are consistent with the results of visualization. It shows that only DTFLOW detects the ICM stage correctly. The intermediate states of cell development in Monocle2 are not revealed properly possibly because the differences between clusters are amplified by the DDRTree method with the cluster centroids. In addition, DPT uses diffusion maps for dimensional reduction, which may not be sensitive enough to the noise in dataset.

We further conduct the robustness analysis of each algorithm. We first use the whole dataset to infer a trajectory and determine the position of each cell in this trajectory. Then we sample part of the cells from the whole dataset and use the same algorithm to determine the trajectory of cells in the sub-dataset. We calculate the Spearman rank correlation coefficient between the positions of subset cells in the trajectory of the whole dataset and those of the sub-dataset. An algorithm is more robust if the value of the correlation coefficient is larger. We conduct 50 tests to measure the robustness properties of these three algorithms. In each test we randomly sample 90% of cells (i.e. 394 cells) from the dataset and then calculate the Spearman rank correlation coefficient of the pseudo-time ordering of the sub-dataset. Then we use the mean and standard deviation of the correlation coefficient based on these 50 test results to measure the robustness property of algorithms. **Figure 3B** shows that the robustness properties of DTFOLW and Scanpy are better than that of Monocle2. In addition, the variance of correlation coefficients obtained by DTFLOW is smaller than that of Scanpy, which suggest that the performance of DTFLOW is more stable than the two published methods. To examine the influence of the sample size, we conduct further robustness test by randomly sampling 80% of cells (i.e. 350 cells) from the dataset. The Spearman rank correlation coefficients in Supplementary Figure S5 are consistent with those shown in **Figure 3B**.

### Mouse myeloid progenitors dataset

We next apply our proposed algorithm to analyze the mouse myeloid progenitor MARS-seq dataset that contains 2730 single cells and 3451 informative genes [41]. Note that 10 genes with corrupted names are removed from our analysis based on the pre-processing of Scanpy. In this experimental study, 19 distinct, transcriptionally homogeneous progenitor types/clusters have been identified through an EM-based clustering approach. Among these clusters, clusters 1∼6 represent Ery (erythroid lineage progenitors) subpopulations, clusters 7∼10 represent CMP (common myeloid progenitors) subpopulations, cluster 11 is for the DC (dendritic cell) fate, clusters 12∼18 correspond to GMP (granulocyte/macrophage progenitors) subpopulations, and cluster 19 is the lymphoid lineage progenitors (outlier class) with only 31 cells. We apply DTFLOW to project this dataset into the two-dimensional feature space. **Figure 4A** elucidates that CMP and its progenitors (namely Ery and GMP) are nearly separated in three different regions, while DC and lymphoid cells deviate away from the main differentiation progression process.

**Figure 4.**
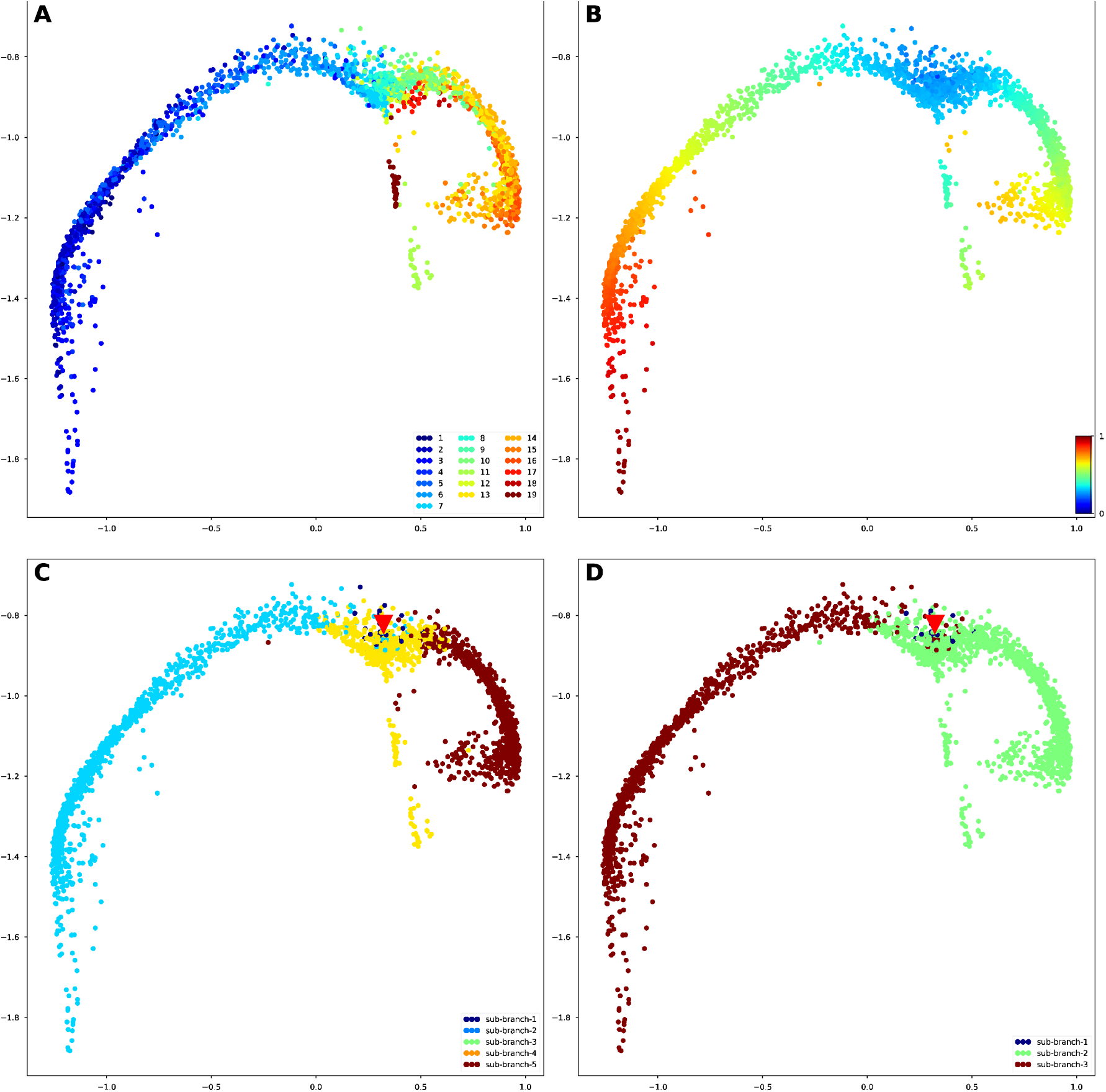
Developmental trajectories inferred by DTFLOW for the mouse myeloid progenitor MARS-seq dataset. A. Visualization of different cell types. B. Visualization of inferred temporal trajectory. C. Visualization of calculated 4 sub-branches when the minimal cell number required for forming a sub-branch satisfies *n =* 6∼22. D. Visualization of calculated 3 sub-branches when the minimal cell number required for forming a sub-branch is larger (*n* = 23∼124).

To reveal the cellular differentiation process, we select the same cell in [31] as the root cell (i.e. the 840-th cell in cluster 8). **Figure 4B** demonstrates the pseudo-time ordering results from the themyeloid progenitor stage. **Figure 4C** and **Figure 4D** show different branching detection results that are determined by a smaller cell number of *n =* 6∼22 and a relatively larger number of *n* = 23∼124 for forming sub-branches, respectively. The first sub-branch in **Figure 4C** contains only a small number of cells. DTFLOW ensures that the pseudo-time of each cell in the initial branch is less than that of any other cells in the following sub-branches. Then cells differentiate into three different terminal branches. Sub-branch two corresponds to the erythroid evolutionary branch, sub-branch three is formed by cells within clusters 11 and 19, and sub-branch four corresponds to the GMP branch. However, when a larger value of *n* is used, sub-branches two and three merge together and form one large sub-branch in **Figure 4D**.

We next compare the branching detection results of DTFLOW, Scanpy and Monocle2. Supplementary Figures S6C and S7C show the branching inference results of Scanpy and Monocle2, respectively. Figure S6C suggests that Scanpy is also able to identify 4 sub-branches. However, the root cell identified by Scanpy is not in the initial group, which is unreasonable for the developmental process. Although Monocle2 successfully estimates 12 states in Figure S7C, which is consistent with the experimental observation, it is difficult to analysis the changes of gene expression over time based on this large branch number.

We then carry out robustness analysis of the three methods. For each method, we randomly sample 2500 single cells from 2730 cells and then use the same methods to infer the pseudo-times of the selected cells. Then we compare the pseudo-times of cells in the sampled set with that of the corresponding cells in the whole dataset by using the Spearman rank correlation coefficient. We conduct 50 repeated tests to measure the robustness property of each method. **Figure 5A** shows that the robustness property of DTFOLW and Scanpy is better than Monocle2. In addition, the variance of correlation coefficients obtained by DTFLOW is smaller than that of Scanpy, which suggest that the performance of DTFLOW is more stable than the two published methods.

**Figure 5.**
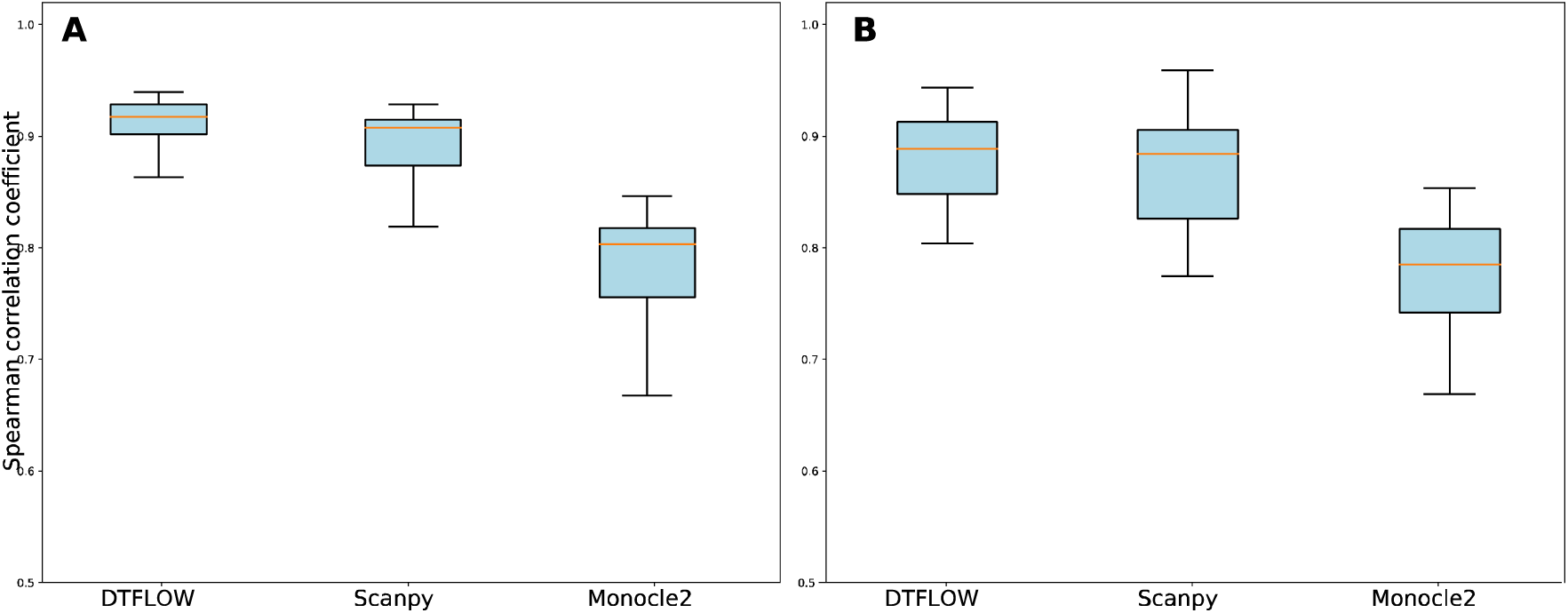
Robustness properties of the three inference methods. A. Robustness property for the mouse myeloid progenitor MARS-seq dataset. B. Robustness property for the mouse female gonad scRNA-seq dataset.

Note that the gene expression levels in this dataset are not continuous and the three terminal branches have different lengths. To illustrate this, Supplementary Figure S8 presents the expression visualization of two marker genes, namely *Elane* and *Klf1*. These marker genes show similar significance in different branches for different dimensionality reduction algorithms. Figure S8A shows that expression levels of gene Elane are essential for the GMP process while gene Klf1 increase gradually on the erythroid branch. The expression trends of these marker genes are different along the constructed trajectories, which provides important information for developing gene regulatory networks.

### Mouse female gonad scRNA-seq dataset

The third dataset is the mouse female gonad scRNA-seq dataset that contains 563 single cells and 822 genes at six developmental stages of gonadal differentiation (namely E10.5, E11.5, E12.5, E13.5, E16.5, and post-natal day 6) [42]. We project this dataset into the 3-dimensional space using our proposed algorithm BKFD. In **Figure 6A** all the cells are presented by different colors for different embryonic stages. It shows that the early progenitor cells subsequently lead to the differentiation to the granulosa cell lineage and stromal progenitor cell lineage in around stages E11.5-E12.5. **Figure 6B** gives the ordered pseudo-times of different single cells and **Figures 6C** presents the inferred three sub-branches by our proposed DTFLOW.

**Figure 6.**
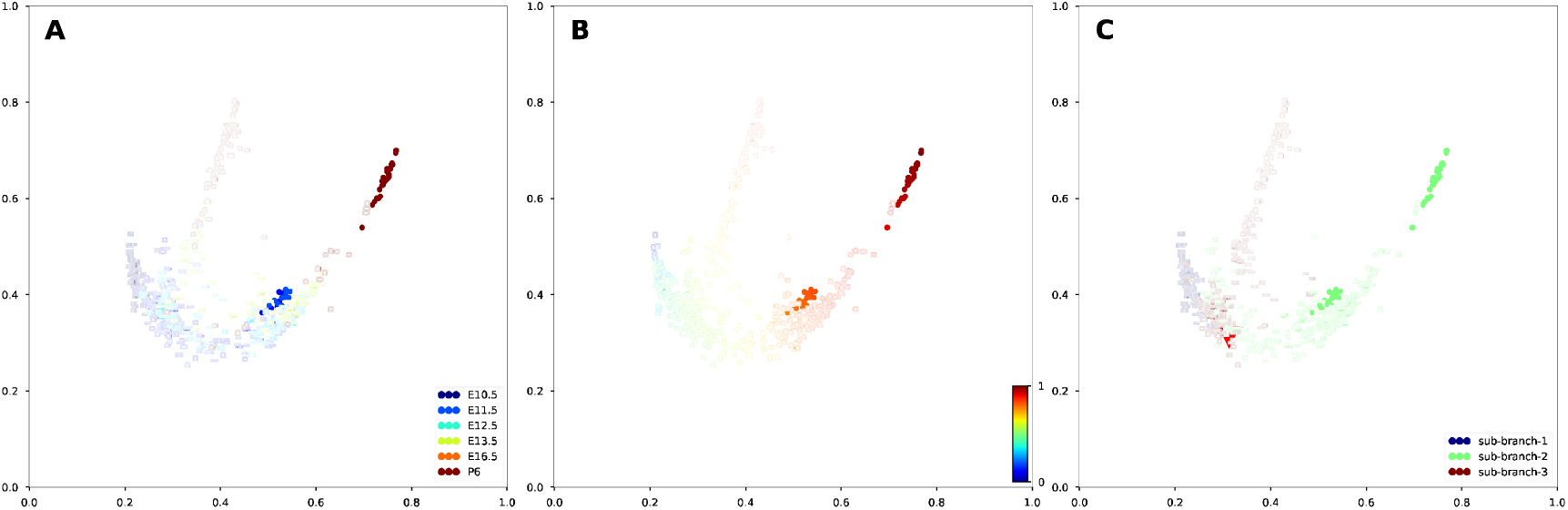
Developmental trajectories inferred by DTFLOW for the mouse female gonad scRNA-seq dataset. A. Visualization of different cell types. B. Visualization of inferred temporal trajectory. C. Visualization of the calculated three sub-branches.

We first compare the pseudo-time ordering accuracy of DTFLOW with Scanpy and Monocle2. Figures S9 and S10 show the analysis results of Scanpy and Monocle2 for this dataset, respectively. The calculated Kendall rank correlations are 0.761, 0.702, 0.569 for DTFLOW, Scanpy and Monocle2, correspondingly. We also compare the robustness properties of DTFOLW with those of Scanpy and Monocle2. We conduct 50 tests to measure the robustness properties of these three algorithms. In each test we randomly sample 90% of cells (i.e. ∼740 cells) from the dataset and find the pseudo-times of these cells. Then we compare the pseudo-times of these cells in the sampled set with that of the corresponding cells in the whole dataset by using the Spearman rank correlation. **Figure 5B** gives the robustness property of these three methods. Numerical results suggest that that the robustness properties of DTFOLW are better than those of Scanpy and Monocle2.

### Comparison of dimensional reduction algorithms

Now we compare our dimensional reduction algorithm BKFD with several popular and widely used methods PCA, tSNE [45] and UMAP [46]. **Figure 7** shows the visualization results of the mouse embryo single-cell dataset [40]. Based on the idea of diffusion propagation, BKFD (**Figure 7A**) represents the cellular development reasonably. **Figure 7B** suggests that PCA cannot distinguish differentiation stages very well. Although tSNE (**Figure 7C**) and UMAP (Figure **Figure 7D**) can separate different cell types clearly for this dataset, the distance intervals of different cells types are relatively large, which cannot be used to indicate cellular developmental processes properly.

**Figure 7.**
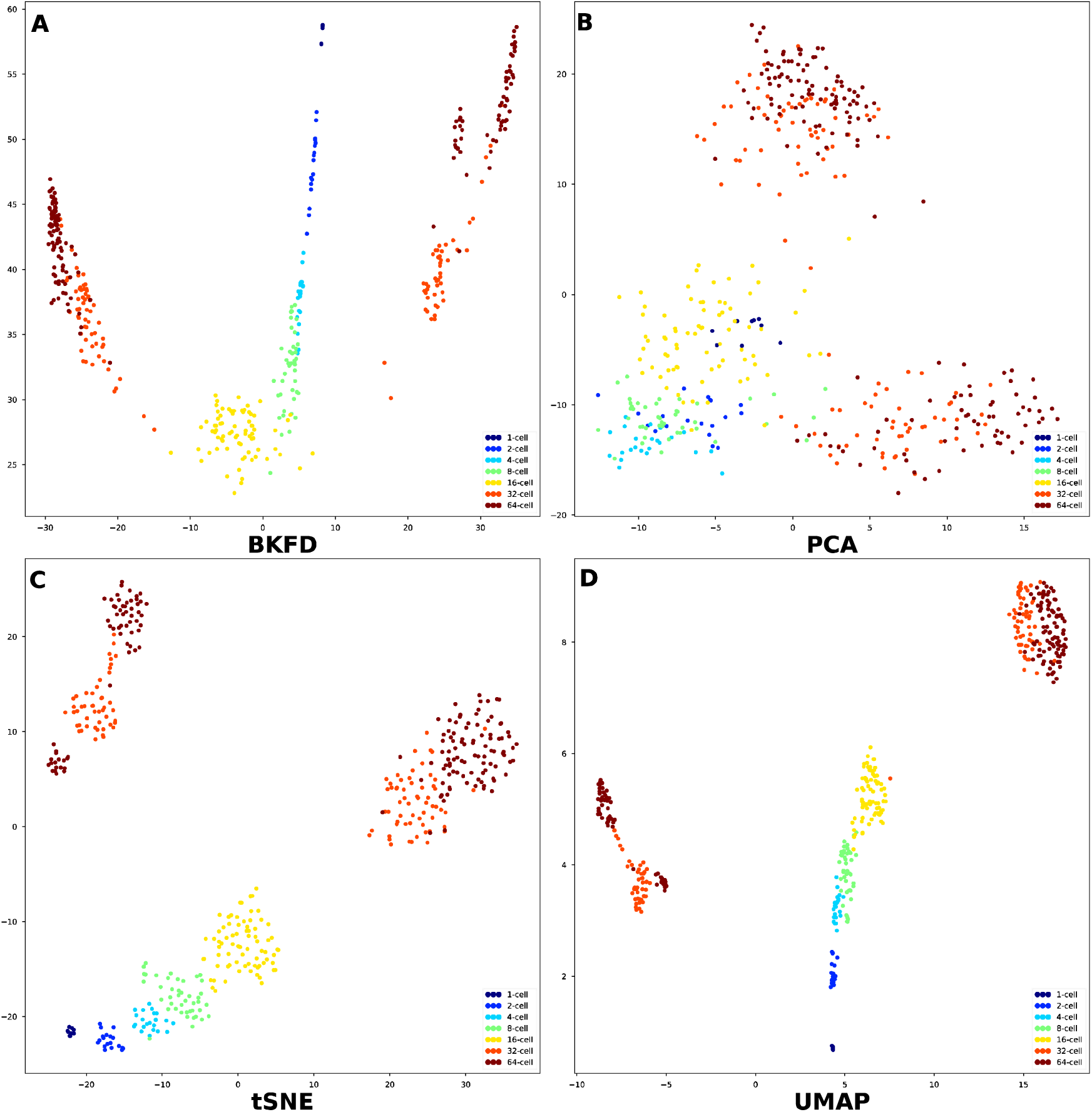
Dimensional reduction results of the dataset [40] by using four different methods. A. BFKD; B.PCA; C. tSNE; D.UMAP.

After the successful applications of our proposed algorithm to three relatively small datasets, we next show the effectiveness of our method to a large dataset. We apply our proposed method BKFD to a microwell-seq data that contains 51252 cells and 25912 genes [43]. After the data pre-processing, the dataset is reduced to 40210 cells with 100 approximate principal components [46]. Based on this dataset, we use four methods for dimensional reduction. **Figure 8** shows the visualization results of this dataset with eight major cell clusters. It suggests that BKFD can capture the developmental trajectories in a better way in **Figure 8A**. In addition, tSNE and UMAP can also distinguish different cell types clearly in **Figures 8C and 8D**, respectively. However, PCA cannot show good visualization results with distinguishable cell clusters for this dataset in **Figure 8B**.

**Figure 8.**
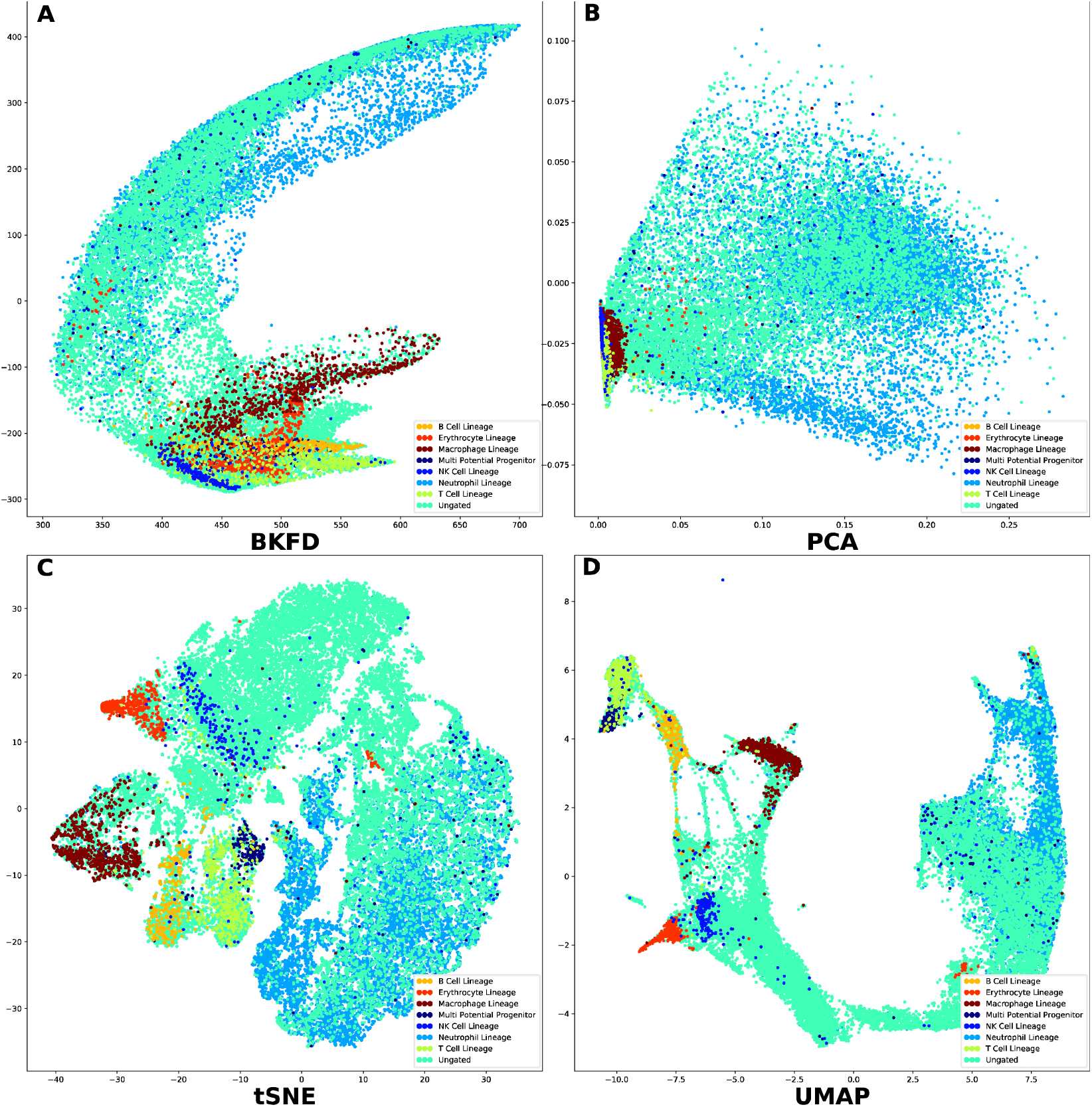
Dimensional reduction results of the dataset [43] by using four different methods. A. BFKD; B.PCA; C. tSNE; D.UMAP.

## Discussion

In this paper we propose a new method DTFLOW for conducting pseudo-time analysis of single-cell data. This method has two major steps: namely a new dimension reduction method BKFD and a novel approach RSKG to identify the underlying multi-branching processes of cellular differentiation. In BKFD we first establish a stationary distribution for each cell to represent the transition of cellular developmental states based on the RWR algorithm, and then propose a new Bhattacharyya kernel matrix to measure the distances between the distributions obtained by RWR. We use this novel distance metric to calculate the pseudo-time distances between single cells before dimension reduction. Thus, our method can reduce the information loss in data processing and increase the inference accuracy. The combination of RWR and the Bhattacharyya kernel matrix shows great power to explore the global structure of the developmental processes using single-cell datasets. In addition, we design the RSKG algorithm to identify the multi-branching of cellular processes. Four datasets are used to compare the accuracy, robustness and branch detection of the proposed algorithm with two popular published methods. Inference results suggest that our proposed method is more accurate and robust than the published algorithms for developing the pseudo-time trajectories of single cells.

The RWR algorithm is a popular method to estimate the global similarity between a particular node point with other node points in the graph structure. We use this method to transform the data of each node point to a stationary discrete distribution. Thus, the input space becomes a set of distributions over the same space. The performance of our proposed algorithm is affected by the choice of Gaussian kernel function, the number of closest neighbours *k* and the restart probability 1 − *p* in the RWR algorithm. Although we have examined the performance of the proposed algorithm by using four datasets, the values of these parameters may vary from dataset to dataset. In addition, BKFD uses the same restart probability for all the nodes, and this may limit the effectiveness of random walk [47]. It is still a challenge to express each cell by a distribution vector in a better way, which needs to be studied in the future.

The continuously topological structure of cellular developmental processes can be analyzed by using the nearest-neighbour graph, which lays the basis of the DTFLOW algorithm. The kNN graph describes the similarities between a cell and its neighbour cells, and has been used twice in the proposed method, namely the definition of transition probability matrix, which leads to the low-dimensional visualization via the Bhattacharyya kernel matrix, and the determination of branching processes in the RSKG algorithm. The new branch detection algorithm identifies the sub-branches through reverse searching on the sequence of indices ordering and provides biological insights into developmental bifurcations. It can ensure that the sub-branches can be connected through the kNN graph, which in turn also verifies its consistency with the pseudo-time inference and visualization results of BKFD.

Scalability is an important issue for the implementation of algorithms. Our algorithm is connected to the dataset size (i.e. the number of cells) in two major steps: the computation of matrix S by finding the inverse of matrix (I-pM) in (6), and the SVD computation in (12). In this study we consider four datasets with cell numbers 438, 2730, 563 and 40210, respectively. The computational time of our algorithm is 0.224 second, 11.65s, 0.246s, and 3108.35s on a Lenovo ThinkPad P53 mobile workstation with 2.6GHz CPU for these four datasets, which is close to the computing time of other algorithms. In addition, the computing time is in the order of *O*(*N*^2^) in terms of the dataset size *N*, which suggests our program is scalable to dataset size.

In summary, the proposed algorithm DTFLOW provides a new framework for inferring the pseudo-time of single cells. Numerical results suggest that it is a power tool for the inference and visualization of cellular developmental trajectories. Potential future work may include the selection of parameters in the proposed method in order to achieve optimal performance in single-cell data analysis.

### Data Accessibility

The first dataset [40] is downloaded from https://github.com/gcyuan/SCUBA/tree/master/sample_data/guo2010. The second dataset[41] is given by https://github.com/theislab/scanpy_usage/tree/master/170502_paul15. The third dataset[42] is download from https://github.com/IStevant/XX-XY-mouse-gonad-scRNA-seq. The fourth dataset [43] is download from https://github.com/ebecht/DR_benchmark.

## Authors’ contributions

JW designed the programs, analysed the data and drafted the manuscript. TZ and XZ analysed and interpreted the data as well as revised manuscript critically. XZ helped programming in revision. TT conceived of the study, participated in the programming and drafting the manuscript. All authors read and approved the final manuscript.

## Acknowledgments

This work was supported by National Natural Science Foundation of China (Grant No. 11571368, 11931019, 11775314, 11871238) and the Fundamental Research Funds for the Central Universities (Grant No. 2662019QD031).

## Supplementary materials

**File S1: Supplementary Information**

## Legends of Supplementary Figures

**Figure S1.**
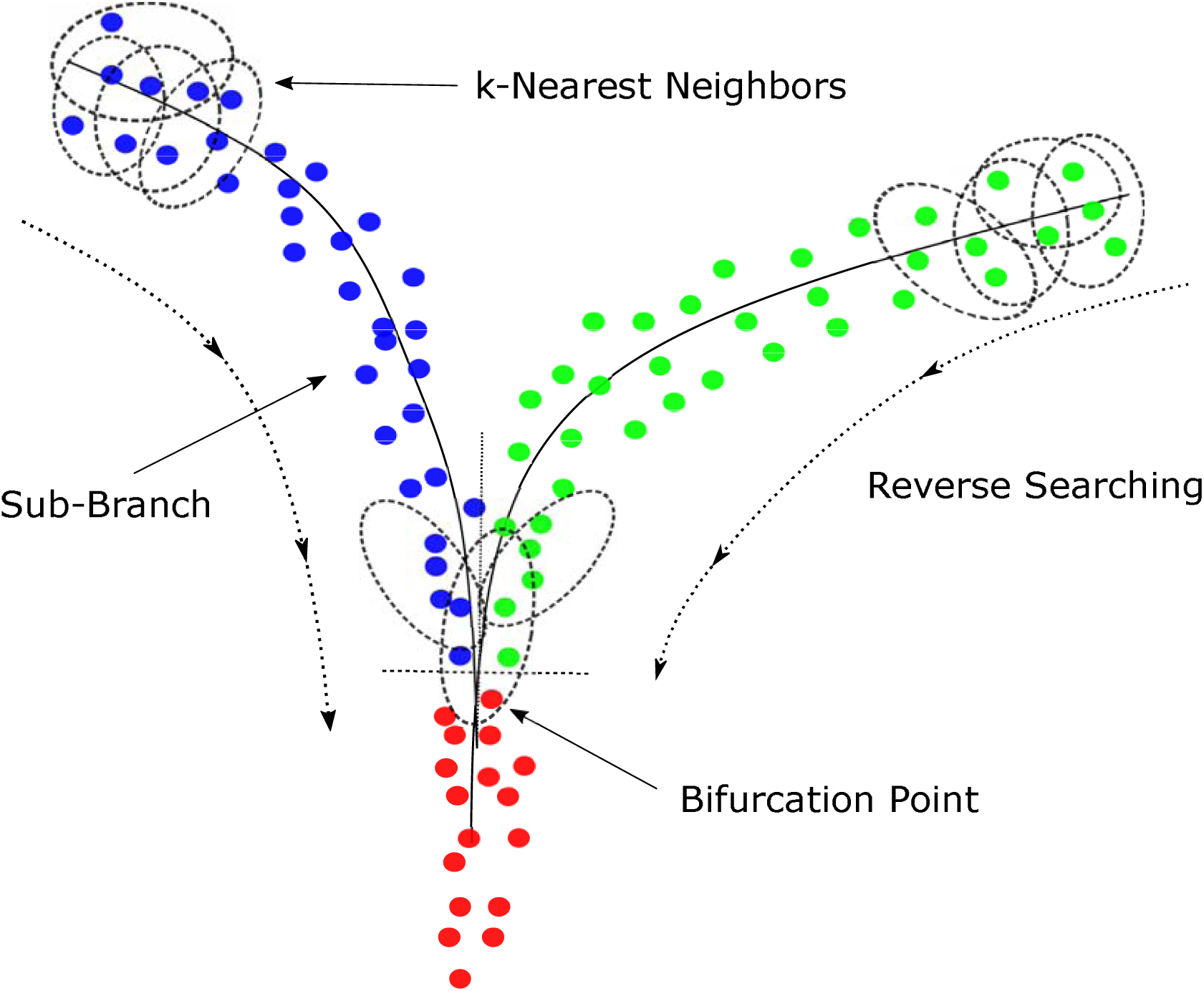
Branch Detection Algorithm by reverse-searching in the kNN Graph.

**Figure S2:**
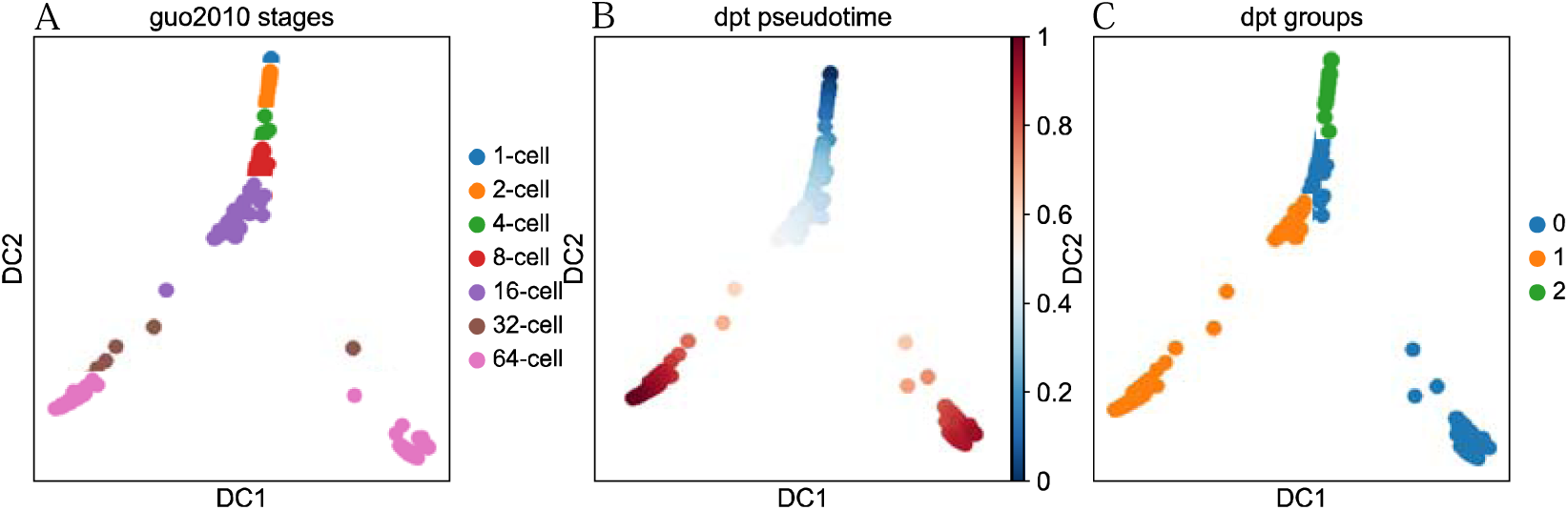
Visualization of the mouse embryo single-cell dataset by using Scanpy. A. Visualization of 7 time stages. B. Visualization of Pseudo-times. C. Visualization of 3 groups/subbranches.

**Figure S3:**
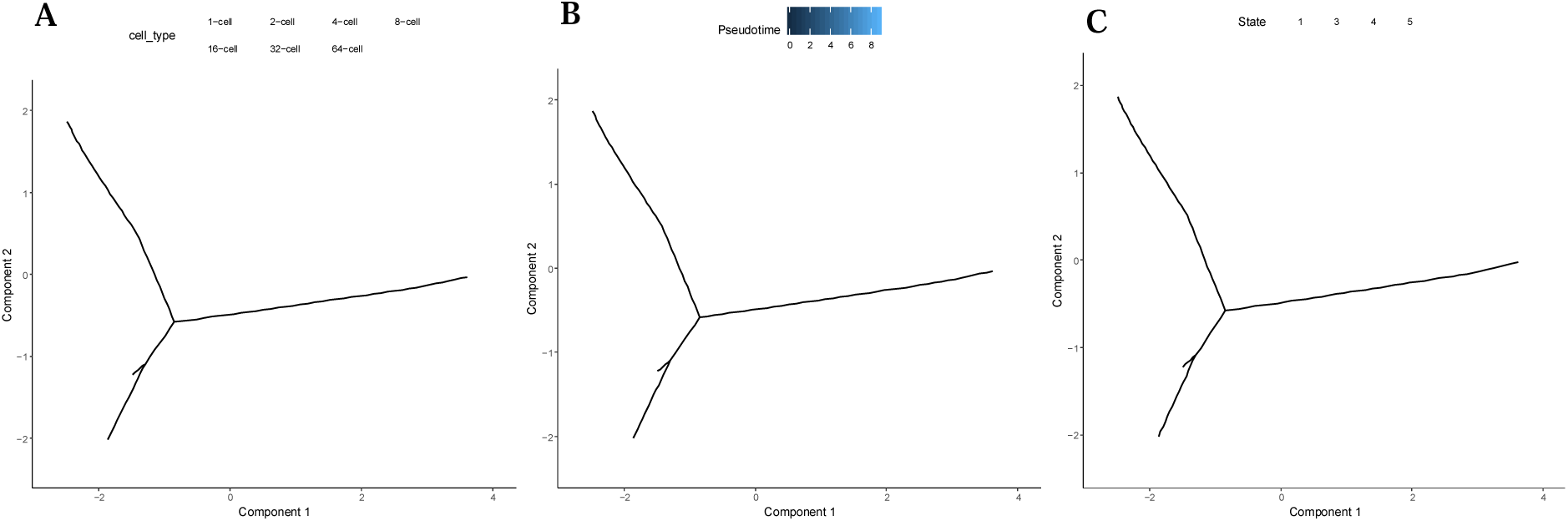
Visualization of the mouse embryo single-cell dataset by using Monocle2. A. Visualization of 7 time stages. B. Visualization of Pseudotimes. C. Visualization of 4 states/sub-branches.

**Figure S4:**
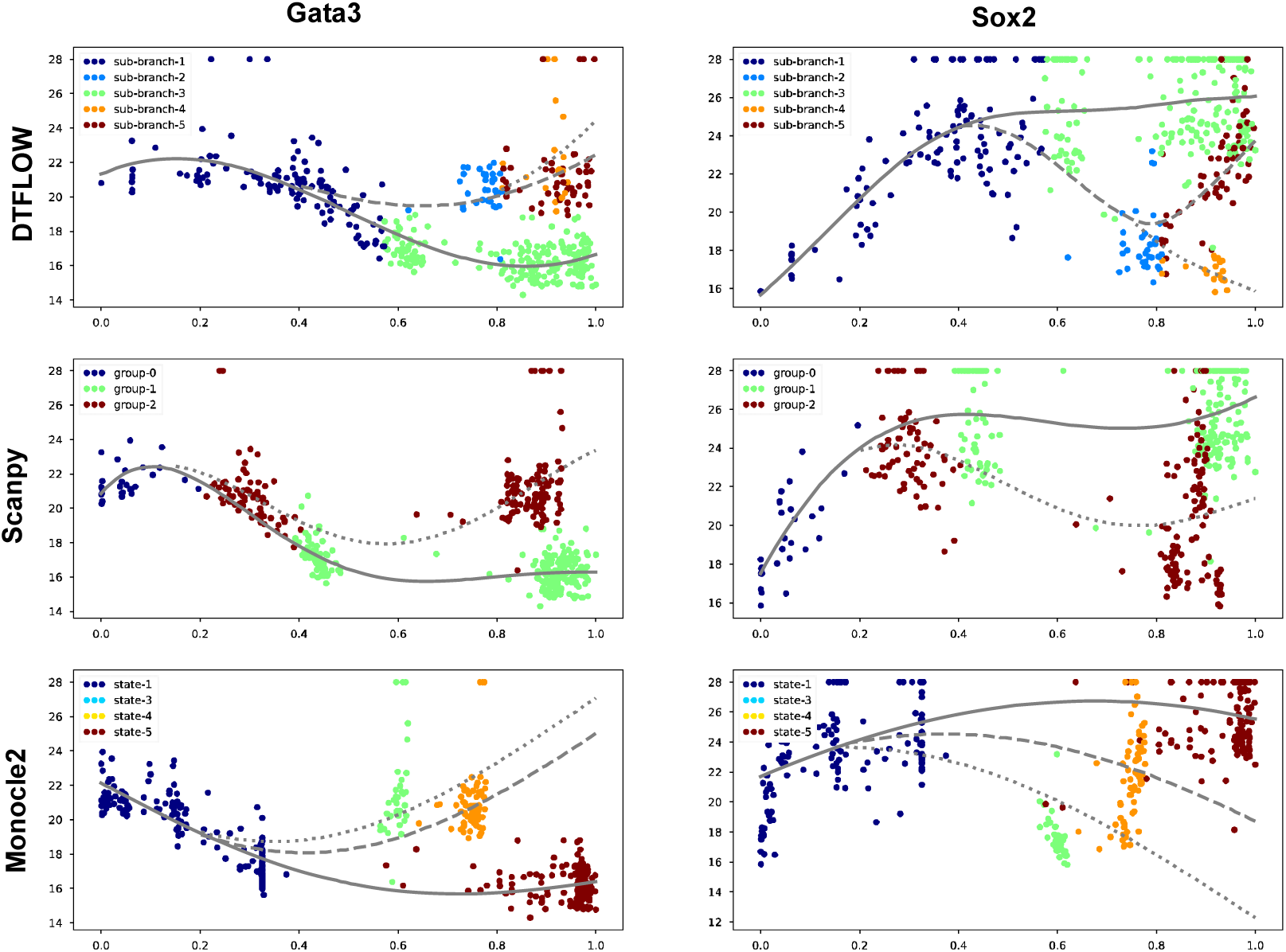
Expression values of genes *Gata3* and *Sox2*. The ordered values from the mouse embryo single-cell dataset are plotted along pseudo-time obtained by using the three inference methods. The lines correspond to the results by using the Gaussion process regression for each branch.

**Figure S5:**
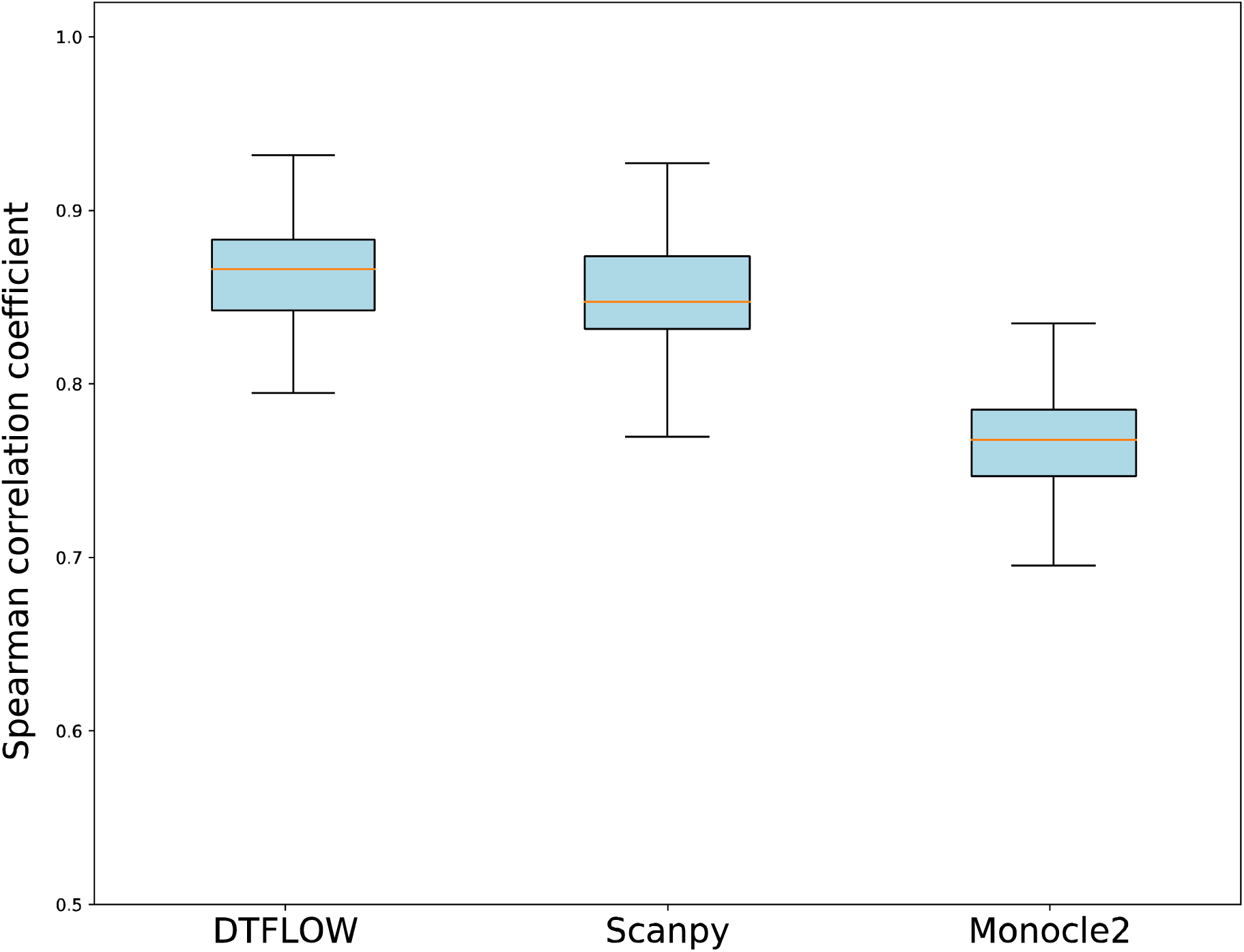
Robustness properties of three inference methods for the mouse embryo single-cell dataset. The properties are obtained by randomly sampling 80% of single-cells from the whole dataset and comparing the ordering results of the subset with those of the whole dataset.

**Figure S6:**
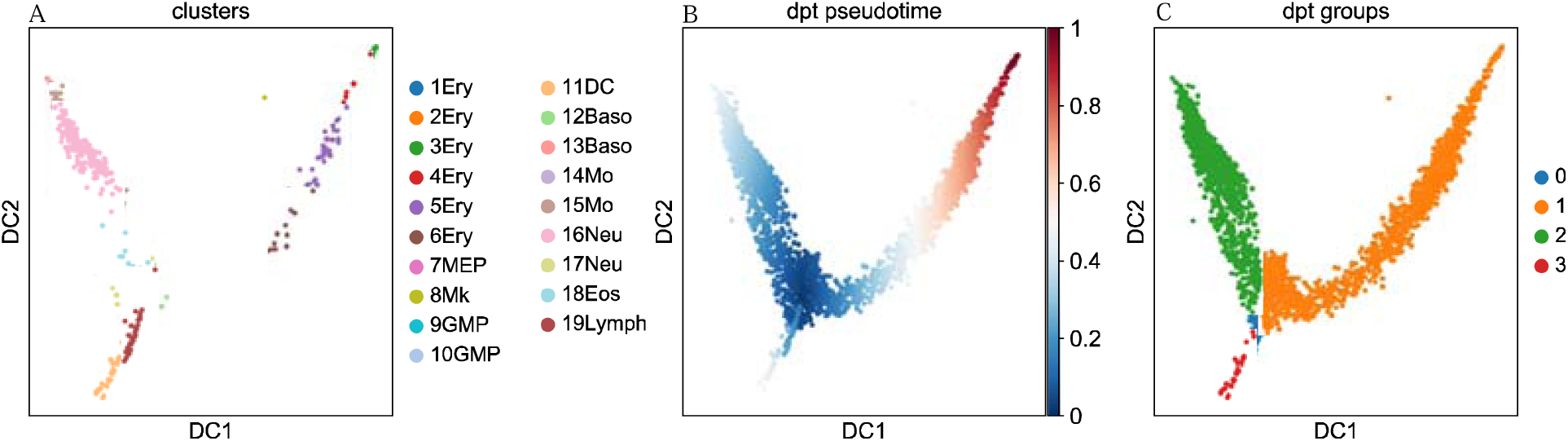
Visualization of the mouse myeloid progenitor MARS-seq dataset by using Scanpy. A. Visualization of 19 cell clusters. B. Visualization of Pseudotimes. C. Visualization of 4 groups/sub-branches.

**Figure S7:**
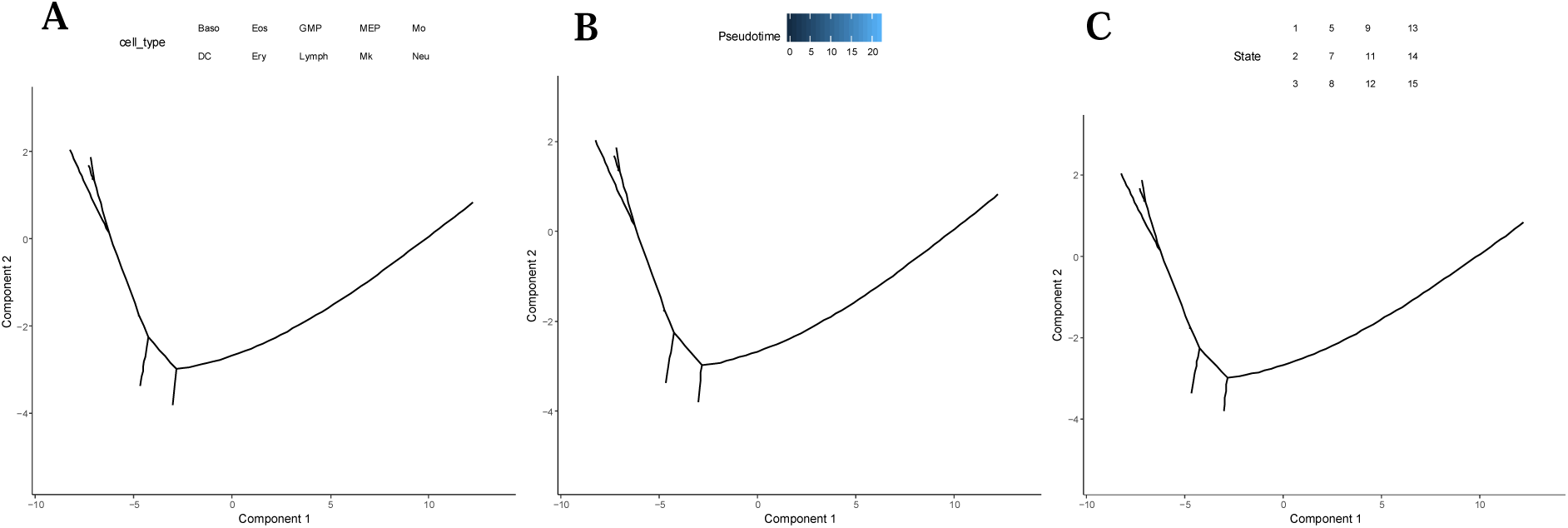
Visualization of the mouse myeloid progenitor MARS-seq dataset by using Monocle2. A.Visualization of 10 cell types. B. Visualization of Pseudo-times. C. Visualization of 12 states/sub-branches.

**Figure S8:**
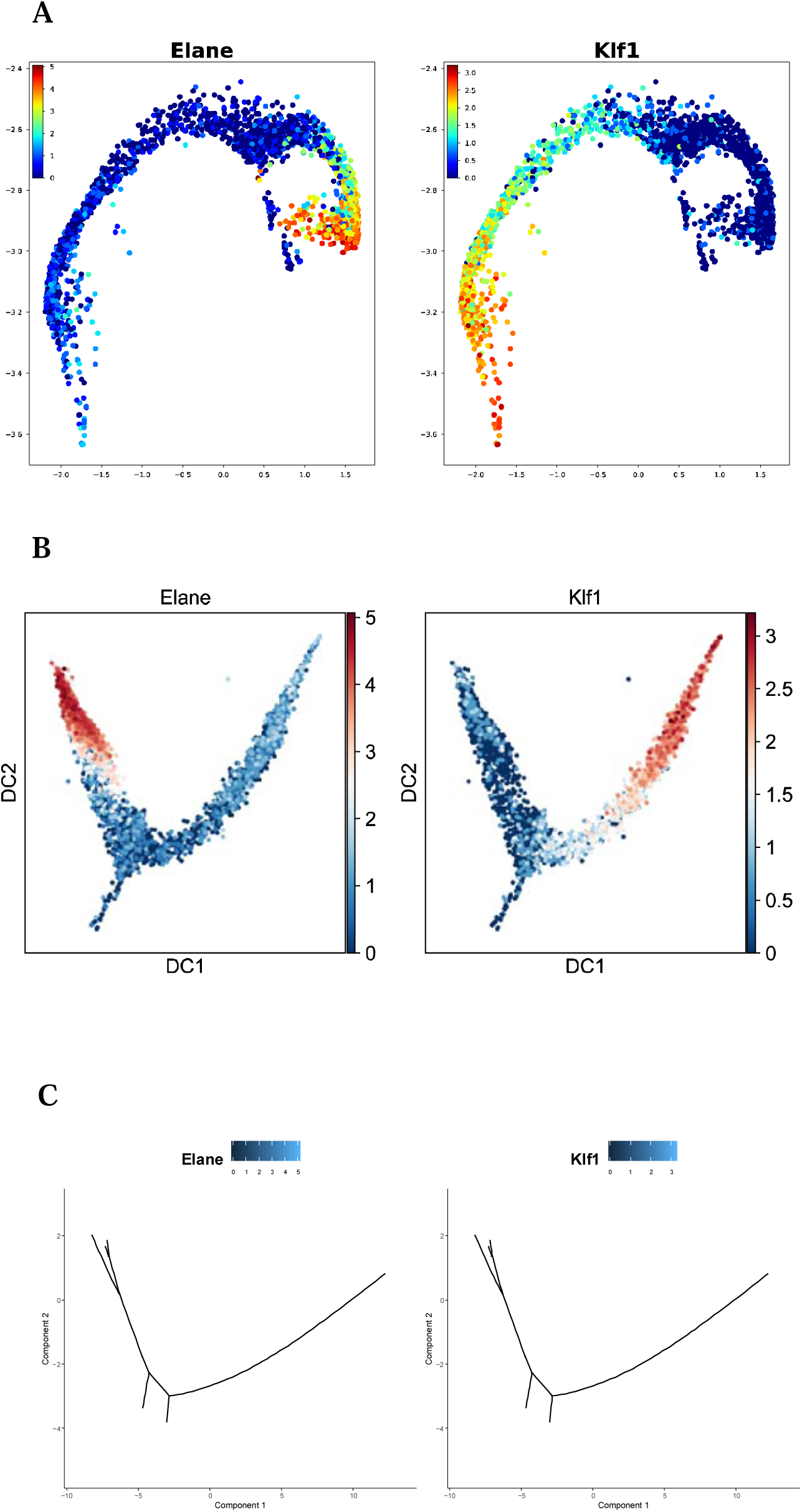
Visualization of gene Elane and Klf1 of the mouse myeloid progenitor MARS-seq dataset by using different methods: A. DTFLOW, B. Scanpy and C. Monocle2.

**Figure S9:**
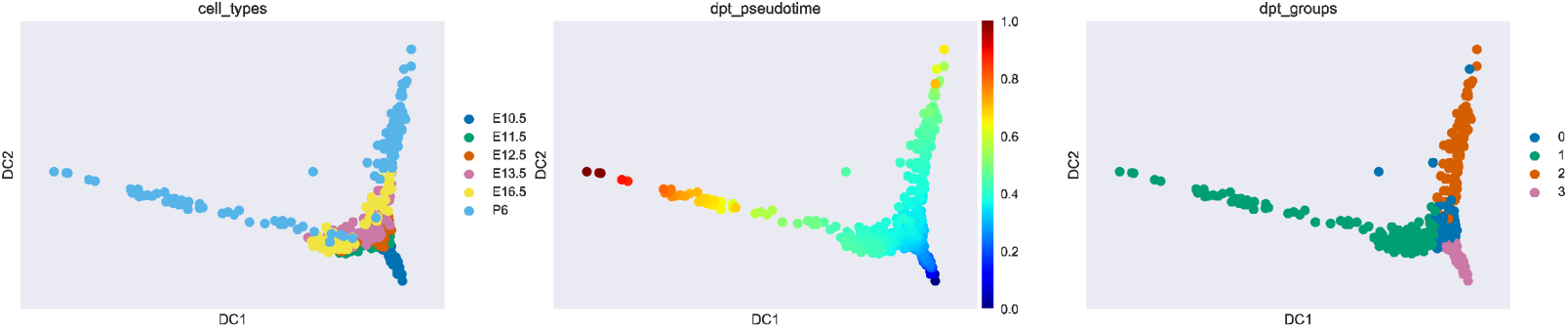
Visualization of the mouse female gonad scRNA-seq dataset by using Scanpy. A. Visualization of 6 cell clusters. B. Visualization of Pseudo-times. C. Visualization of 4 groups/sub-branches.

**Figure S9:**
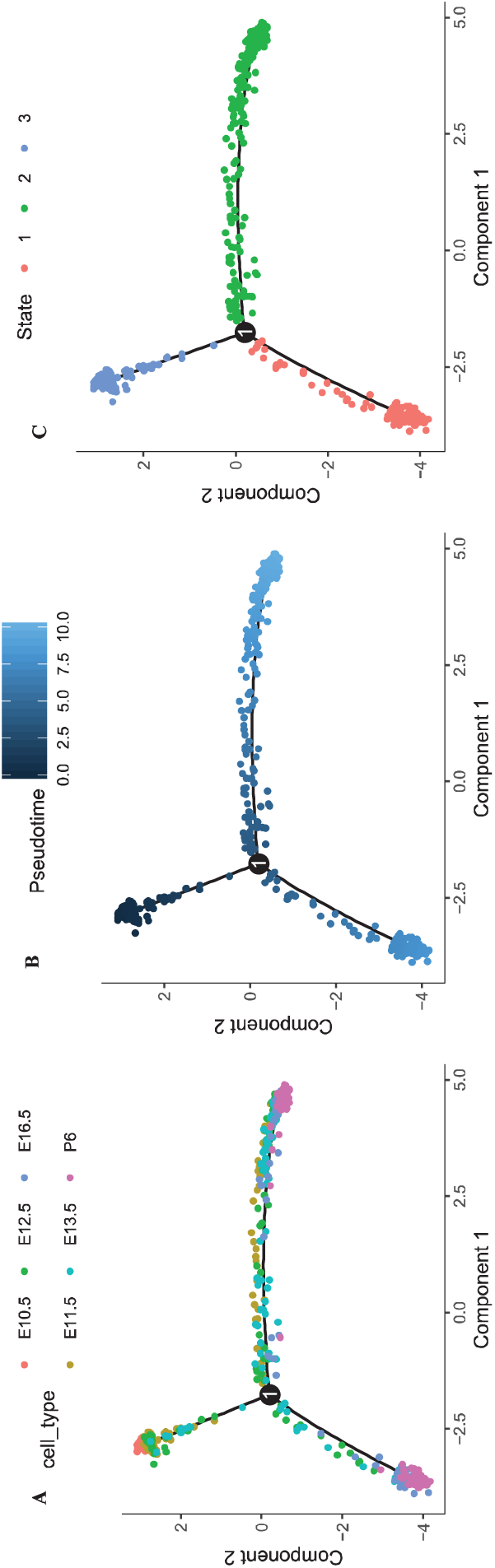
Visualization of the mouse female gonad scRNA-seq dataset by using Monocle2. A. Visualization of 6 cell types. B. Visualization of Pseudo-times. C. Visualization of 3 states/sub-branches.

## Supplementary Information

### S1. Comparison between DPT and DTFLOW

We note that DPT and DTFLOW use the similar propagation process. DPT provides the (time independent) “path integral” by

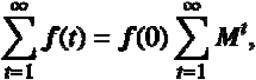

where is a probability density. Then the diffusion process of DPT is given by

where the eigenvector is corresponding to the eigenvalue 1 of the transition matrix. The diffusion pseudo-time distance metric *dpt* is given by

which implies that it is also a kernel distance based on one kernel matrix.

For the DTFLOW algorithm, the equation (7) can also be written as

Thus, DTFLOW can use parameter to control the propagation procedure, which may be the reason that the accuracy of DTFLOW is better than DPT.

### S2. The DTFLOW algorithm

Algorithm 1 below gives the detailed steps of DTFLOW.

#### Algorithm 1. DTFLOW

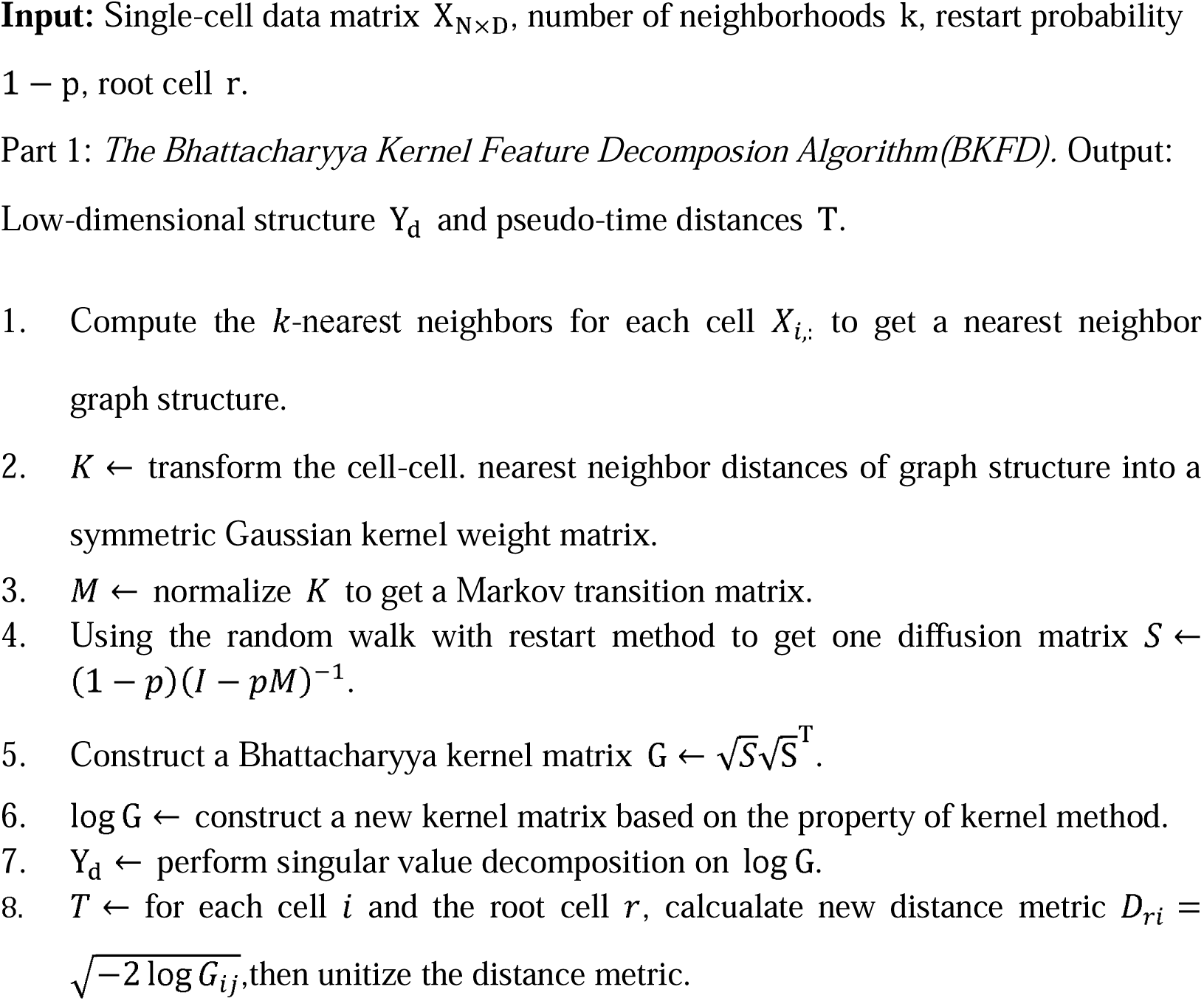

Part 2. Branch Detection Algorithm by reverse-searching in the kNN Graph (see Algorithm 2).

### S3. Branch Detection Algorithm by reverse-searching in the kNN Graph

#### Algorithm 2. RSKG

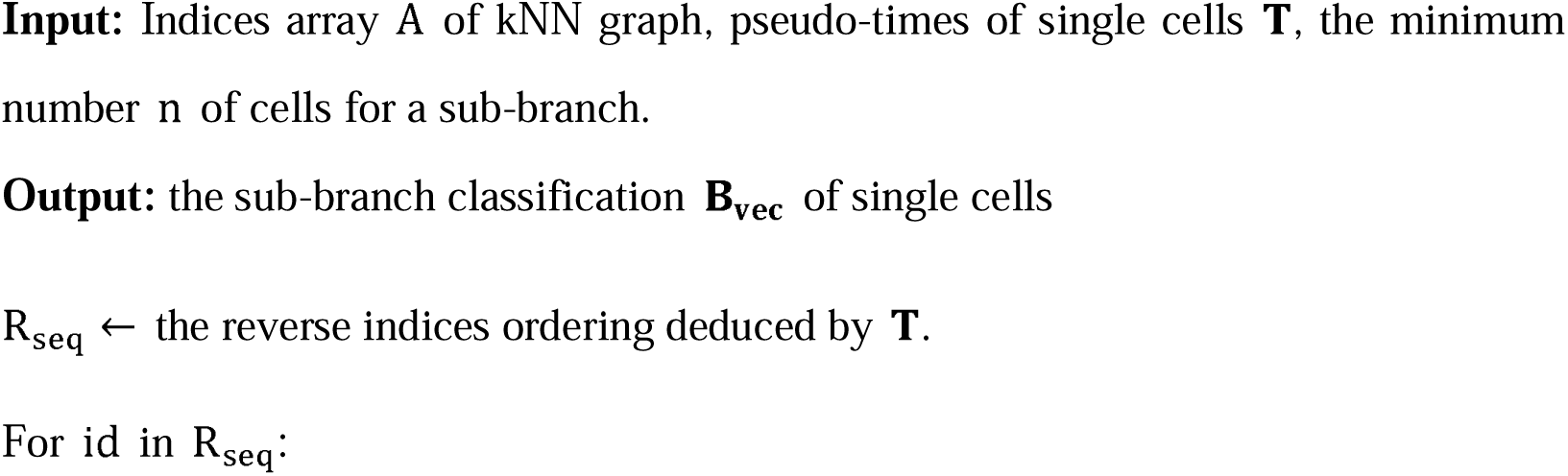
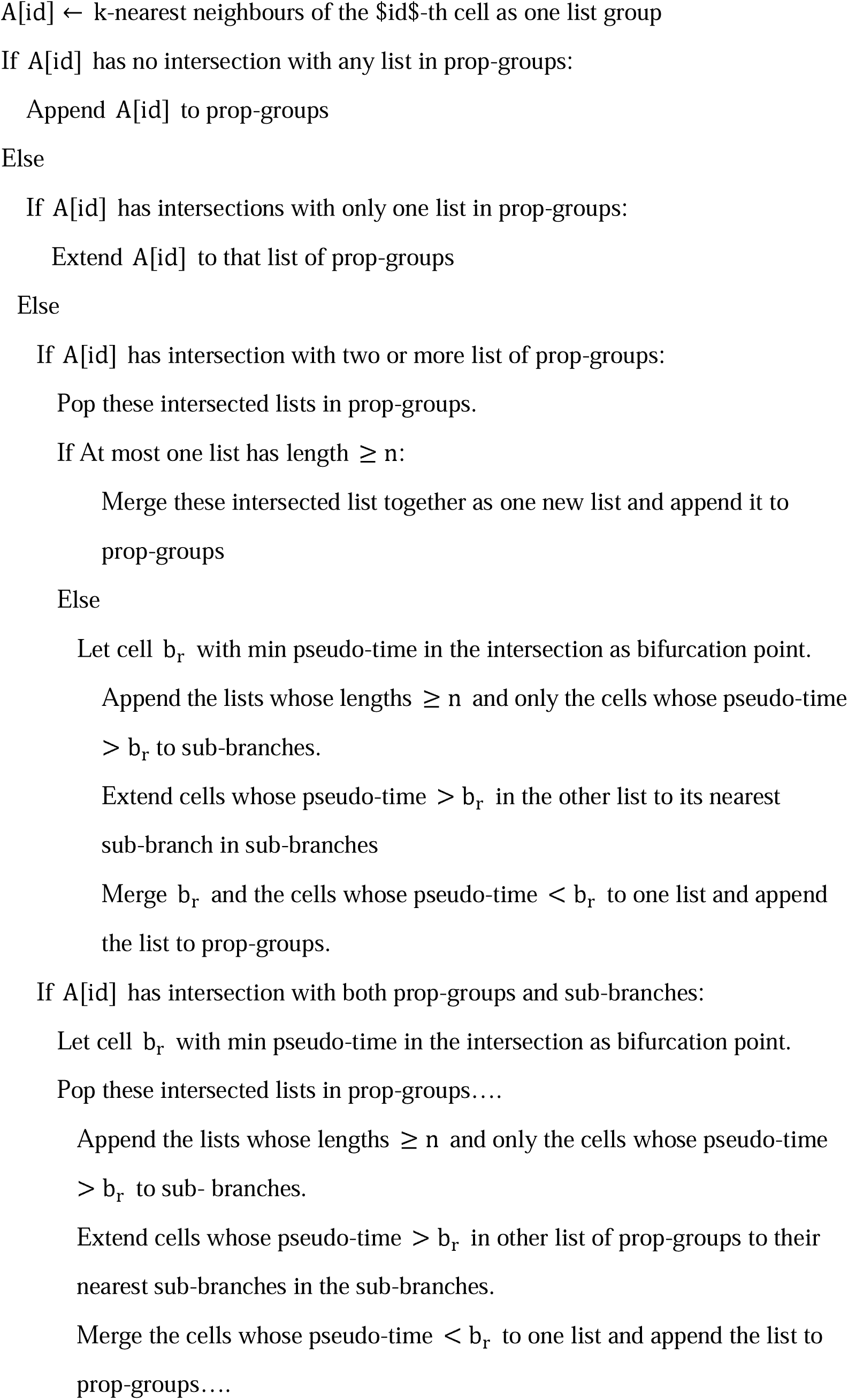

Assign the remaining lists in prop-groups to sub-branches, then get B_vec_ based on sub-branches.

