## Supplementary Information for "DTFLOW: Inference and Visualization of Single-cell Pseudo-temporal Trajectories Using Diffusion Propagation"

**S1. Comparison between DPT and DTFLOW**

We note that DPT and DTFLOW use the similar propagation process. DPT provides the (time independent) "path integral" by

$$\sum_{t=1}^{\infty} f\left( t \right)=f\left( 0 \right)\sum_{t=1}^{\infty} M^{t},$$

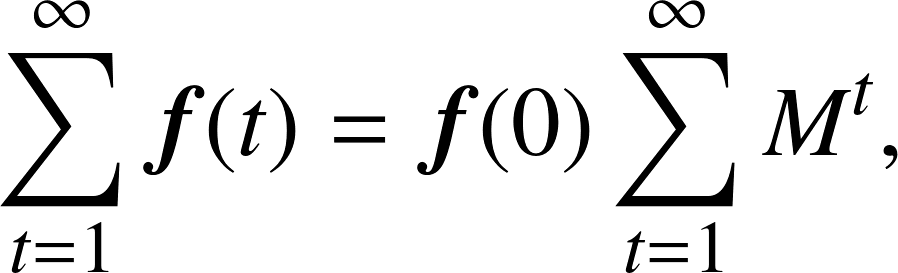

where $f\left( t \right)\in R^{n}$ is a probability density. Then the diffusion process of DPT is given by

$$S=\sum_{k=1}^{\infty} \left( M-\psi_{0}\psi_{0}^{T} \right)^{k}=\left( I-\left( M-\psi_{0}\psi_{0}^{T} \right) \right)^{-1}-I,$$

where the eigenvector $\psi_{0}$ is corresponding to the eigenvalue 1 of the transition matrix $M$. The diffusion pseudo-time distance metric *dpt* is given by

$$dpt^{2}\left( x_{i},x_{j} \right)=\left| \left| S\left( x_{i},\cdot\right)-S\left( x_{j},\cdot\right) \right| \right|^{2},$$

which implies that it is also a kernel distance based on one kernel matrix $SS^{T}$.

For the DTFLOW algorithm, the equation (7) can also be written as

$$S=\left( 1-p \right)\left( I+pM+p^{2}M^{2}+\cdots+p^{t}M^{t}+\cdots\right)$$

$$=\left( 1-p \right)\left( I+\sum_{t=1}^{\infty} p^{t}M^{t} \right) =\left( 1-p \right)\left( I-pM \right)^{-1}.$$

Thus, DTFLOW can use parameter $p$ to control the propagation procedure, which may be the reason that the accuracy of DTFLOW is better than DPT.

**S2. The DTFLOW algorithm**

Algorithm 1 below gives the detailed steps of DTFLOW.

**Algorithm 1. DTFLOW**

**Input:** Single-cell data matrix $X_{N\times D}$, number of neighborhoods $k$, restart probability $1-p$, root cell $r$.

Part 1: The Bhattacharyya Kernel Feature Decomposion Algorithm(BKFD). Output: Low-dimensional structure $Y_{d}$ and pseudo-time distances $T$.

1. Compute the $k\text{-nearest}$ neighbors for each cell $X_{i,:}$ to get a nearest neighbor graph structure.
2. $K\leftarrow$ transform the cell-cell. nearest neighbor distances of graph structure into a symmetric Gaussian kernel weight matrix.
3. $M\leftarrow$ normalize $K$ to get a Markov transition matrix.
4. Using the random walk with restart method to get one diffusion matrix $S\leftarrow\left( 1-p \right)\left( I-pM \right)^{-1}$.
5. Construct a Bhattacharyya kernel matrix $G\leftarrow\sqrt{S}\sqrt{S}^{T}$.
6. $\log G\leftarrow$ construct a new kernel matrix based on the property of kernel method.
7. $Y_{d}\leftarrow$ perform singular value decomposition on $\log G$.
8. $T\leftarrow$ for each cell $i$ and the root cell $r$, calcualate new distance metric $D_{ri}=\sqrt{-2\log G_{ij}}$,then unitize the distance metric.

Part 2. Branch Detection Algorithm by reverse-searching in the kNN Graph (see Algorithm 2).

**S3. Branch Detection Algorithm by reverse-searching in the kNN Graph**

**Algorithm 2. RSKG**

**Input:** Indices array $A$ of kNN graph, pseudo-times of single cells $\mathbf{T}$, the minimum number $n$ of cells for a sub-branch.

**Output:** the sub-branch classification $\mathbf{B}_{\mathbf{vec}}$ of single cells

$R_{\mathrm{seq}}\leftarrow$ the reverse indices ordering deduced by $\mathbf{T}$.

For $\mathrm{id}$ in $R_{\mathrm{seq}}$:

$A\left[ \mathrm{id} \right]\leftarrow$ $k$-nearest neighbours of the $id$-th cell as one list group

If $A[id]$ has no intersection with any list in prop-groups:

Append $A[id]$ to prop-groups

Else

If $A\left[ \mathrm{id} \right]$ has intersections with only one list in prop-groups:

Extend $A[id]$ to that list of prop-groups

Else

If $A[id]$ has intersection with two or more list of prop-groups:

Pop these intersected lists in prop-groups.

If At most one list has length $\geq n$:

Merge these intersected list together as one new list and append it to prop-groups

Else

Let cell $b_{r}$ with min pseudo-time in the intersection as bifurcation point.

Append the lists whose lengths $\geq n$ and only the cells whose pseudo-time $>b_{r}$to sub-branches.

Extend cells whose pseudo-time $>b_{r}$ in the other list to its nearest sub-branch in sub-branches

Merge $b_{r}$ and the cells whose pseudo-time $<b_{r}$ to one list and append the list to prop-groups.

If $A[id]$ has intersection with both prop-groups and sub-branches:

Let cell $b_{r}$ with min pseudo-time in the intersection as bifurcation point.

Pop these intersected lists in prop-groups….

Append the lists whose lengths $\geq n$ and only the cells whose pseudo-time $>b_{r}$ to sub- branches.

Extend cells whose pseudo-time $>b_{r}$ in other list of prop-groups to their nearest sub-branches in the sub-branches.

Merge the cells whose pseudo-time $<b_{r}$ to one list and append the list to prop-groups….

Assign the remaining lists in prop-groups to sub-branches, then get $B_{\mathrm{vec}}$ based on sub-branches.

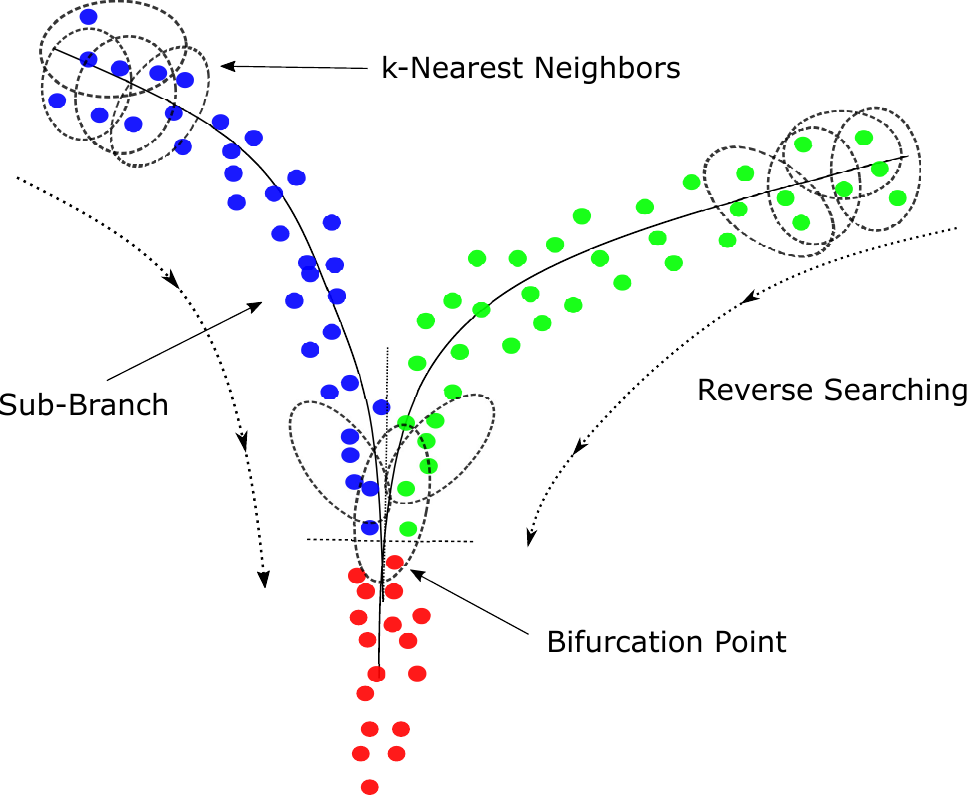

Supplementary Figure 1.

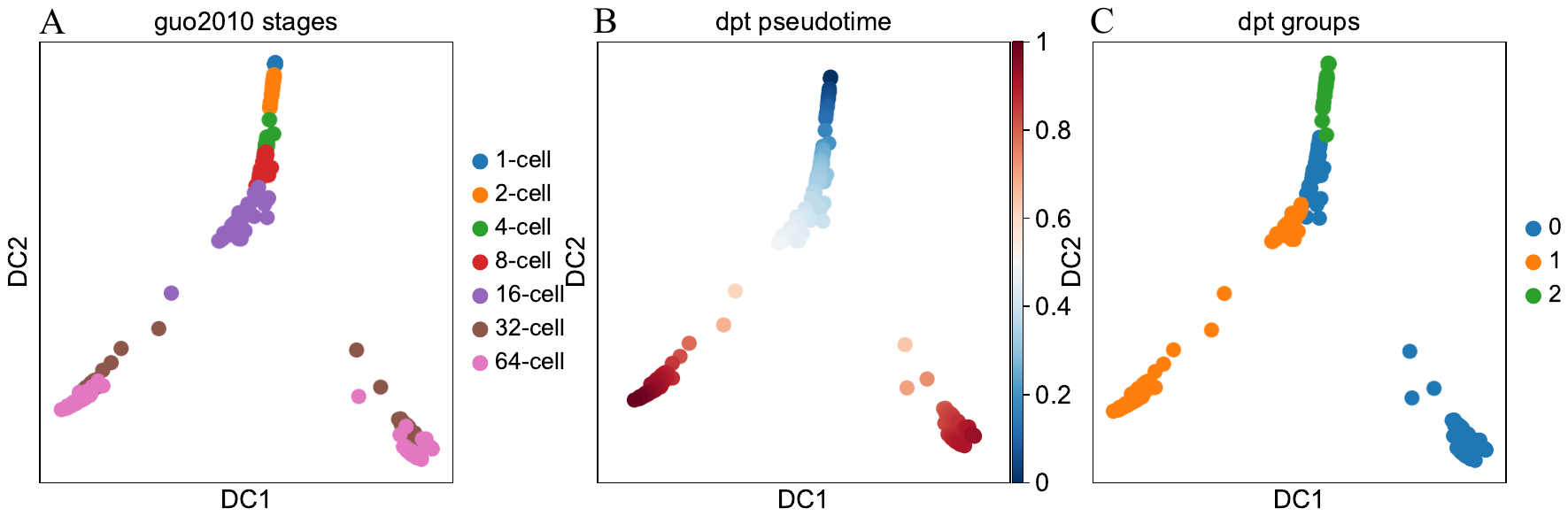

Supplementary Figure 2.

Supplementary Figure 3.

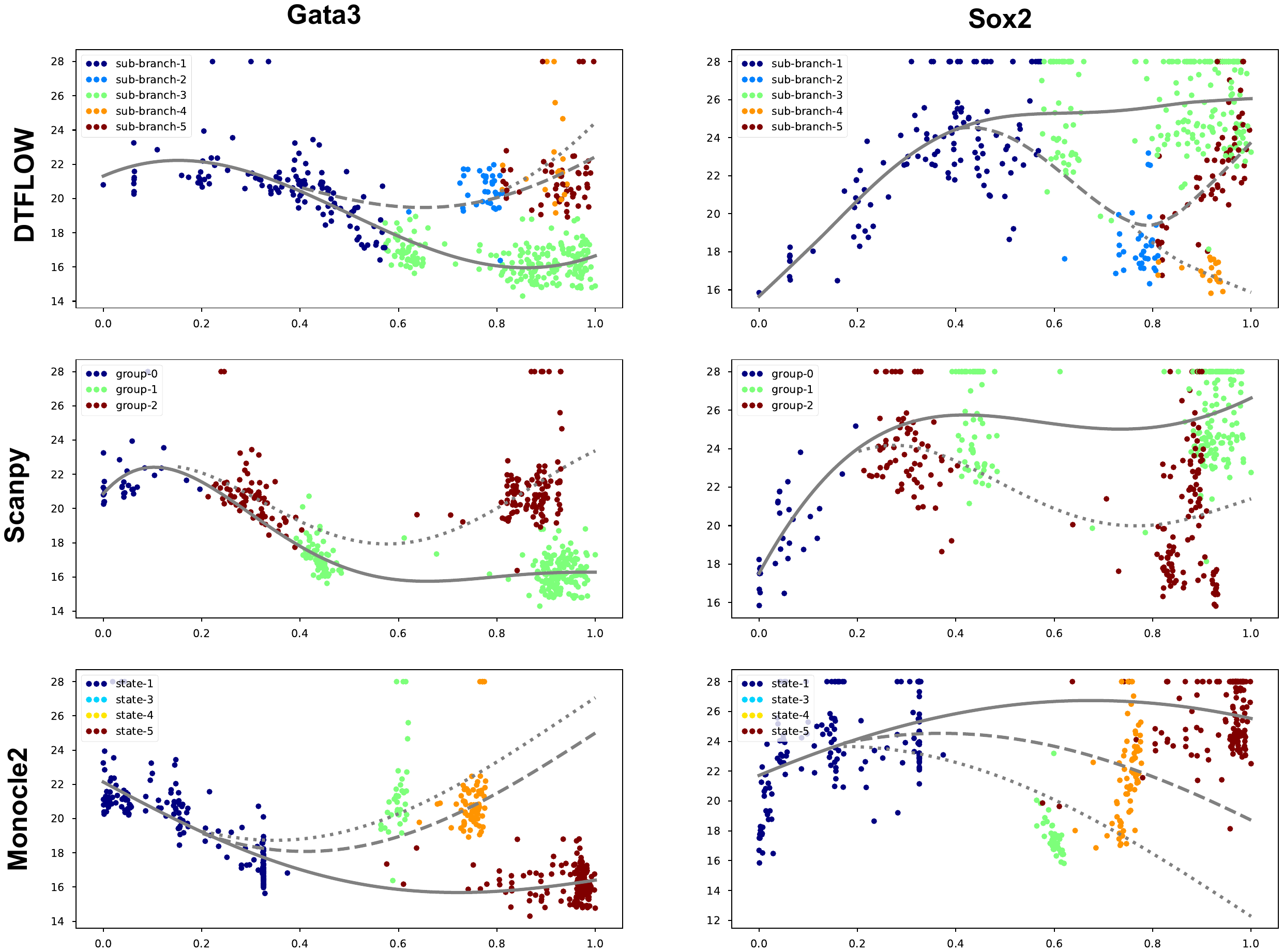

Supplementary Figure 4.

Supplementary Figure 5.

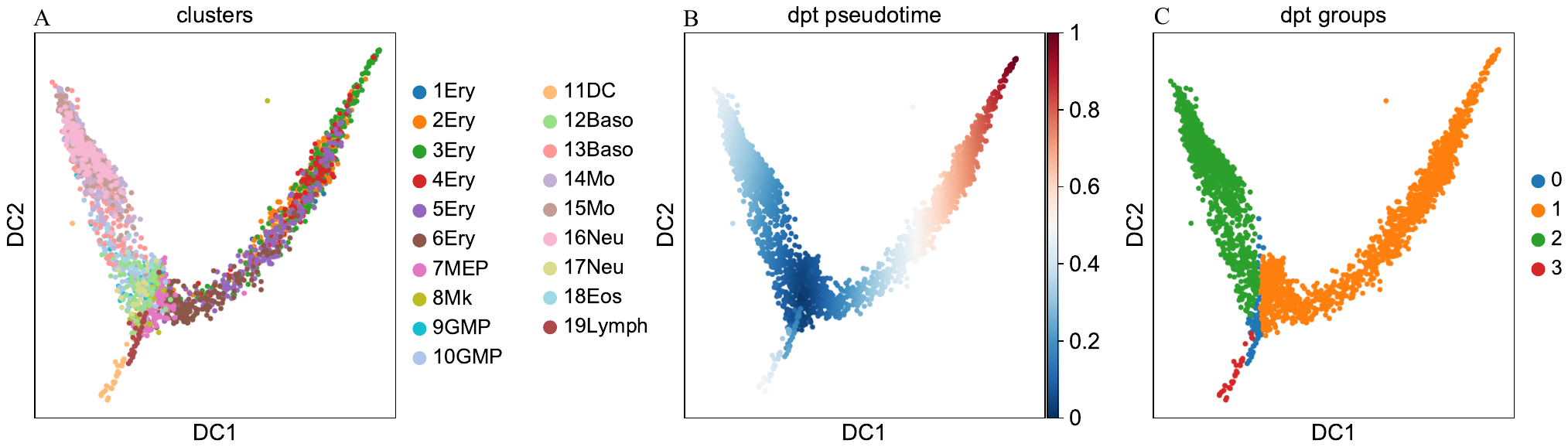

Supplementary Figure 6.

Supplementary Figure 7.

Supplementary Figure 8.

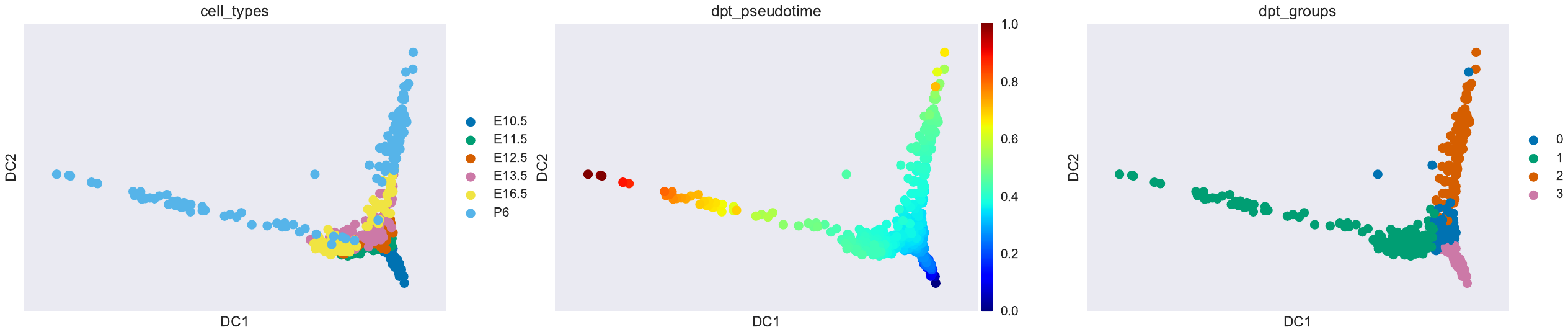

Supplementary Figure 9.

Supplementary Figure 10.
